# Comprehensive analysis of chemical and biological problems associated with browning agents used in aquatic studies

**DOI:** 10.1101/2021.02.26.433092

**Authors:** Kristin Scharnweber, Sari Peura, Katrin Attermeyer, Stefan Bertilsson, Lucas Bolender, Moritz Buck, Karólína Einarsdóttir, Sarahi L. Garcia, Raphael Gollnisch, Charlotte Grasset, Marloes Groeneveld, Jeffrey A. Hawkes, Eva S. Lindström, Christin Manthey, Robyn Övergaard, Karin Rengefors, Vicente T. Sedano-Núñez, Lars J. Tranvik, Anna J. Székely

**Affiliations:** Uppsala University, Department of Ecology and Genetics, Limnology, Norbyvägen 18D, 7523618 Uppsala, Sweden; Swedish University of Agricultural Sciences, Department of Forest Mycology and Plant Pathology, Science for Life Laboratory, Almas allé 5, 75651 Uppsala, Sweden; WasserCluster Lunz, Dr. Carl Kupelwieser Promenade 5, 3293 Lunz am See, Austria; Lund Swedish University of Agricultural Sciences, Department of Aquatic Sciences and Assessment, Lennart Hjelms väg 9, SE75651 Uppsala, Sweden; Stockholm University, Department of Ecology, Environment, and Plant Sciences, Science for Life Laboratory, Svante Arrhenius väg 20 A, 114 18 Stockholm, Sweden; Lund University, Department of Biology, Sölvegatan 37, 22362 Lund, Sweden; Uppsala University, Analytical Chemistry, Department of Chemistry BMC, Husargatan 3, 75237 Uppsala, Sweden; Freie Universität Berlin, Institute of Biology, Evolutionary Biology, Königin-Luise-Str. 1-3, 14195 Berlin-Dahlem, Germany; Uppsala University, Molecular Evolution, Department of Cell and Molecular Biology BMC, Husargatan 3, 75237 Uppsala, Sweden

**Keywords:** HuminFeed, SuperHume, dissolved organic carbon, humic substances, Daphnia, microbial community, *Gonyostomum semen*, terrestrial carbon, reverse osmosis, leonardite

## Abstract

Inland waters receive and process large amounts of colored organic matter from the terrestrial surroundings. These inputs dramatically affect the chemical, physical, and biological properties of water bodies, as well as their roles as global carbon sinks and sources. To understand the complex changes associated with allochthonous inputs, experiments are needed. However, manipulative studies, especially at ecosystem scales, require large amounts of dissolved organic matter with optical and chemical properties resembling indigenous organic matter. Here we compared the chemical and biological impacts of two leonardite products (HuminFeed (HF) and SuperHume (SH)) and a freshly derived reverse osmosis concentrate of organic matter (RO) in a set of comprehensive mesocosm- and laboratory-scale experiments and analyses.

The chemical properties of RO concentrate and the leonardite products were very different with leonardite products being low and RO being high in carboxylic functional groups. Light had a strong impact on the properties of leonardite products, including loss of color and increased particle formation. Furthermore, HF had drastic impacts on bacteria as light stimulated bacterial production and modified community composition, while dark conditions appeared to inhibit bacterial processes. While none of the browning agents inhibited the growth of the tested phytoplankton, *Gonyostomum semen*, leonardite products had detrimental effects on zooplankton abundance and *Daphnia* reproduction. We conclude that the effects of browning agents extracted from leonardite are in sharp contrast to those originating from terrestrially-derived DOM. Hence, they should be used with great caution in experimental studies on the consequences of terrestrial carbon for aquatic systems.

## Introduction

Inland waters process large amounts of terrestrial organic carbon (Cole, Prairie, Caraco and others 2007; Drake, Raymond and Spencer 2018; Tranvik, Cole and Prairie 2018). In the last decades, an increasing load of terrestrially derived dissolved organic matter (DOM) in aquatic systems of the Northern hemisphere, known as “browning”, has been described (Monteith, Stoddard, Evans and others 2007; Solomon, Jones, Weidel and others 2015). Browning has diverse consequences for aquatic ecosystems, largely due to more efficient absorption of solar radiation that alters the vertical distribution of heat and light (Fee, Hecky, Kasian and others 1996; Kirk 2011). This leads to cooler deep waters while the shading also hampers photosynthesis, and thereby reduces algal food supply for higher trophic levels such as zooplankton or fish (Kelly, Solomon, Weidel and others 2014). All these mechanisms influence vertical habitat gradients, food web structures, resource subsidies, and ultimately, ecosystem services (Williamson, Overholt, Pilla and others 2015). Thus, as browning has a high potential to affect ecosystem functioning and water quality, as well as to further aggravate greenhouse gas emissions, it has become a primary subject of experimental studies targeting climate change impacts on freshwaters (Bergström and Karlsson 2019; Vasconcelos, Diehl, Rodriguez and others 2019; Weyhenmeyer, Müller, Norman and others 2016).

One challenge of experimental studies of browning is to find a browning agent that can be applied at different experimental scales and ideally also enables disentangling the impact of increasing organic carbon substrates from the impact of physical darkening of the water column. Browning agents previously applied include extracts of humic substances from soils (e.g. Lennon and Cottingham 2008), leachates from organic material (e.g. Geddes 2009), or the use of DOC-rich waters (e.g. Kritzberg, Graneli, Bjork and others 2014). However, obtaining sufficient quantities of such materials to enable experimental manipulation at mesocosm or ecosystem scale, is challenging and time consuming. A further challenge is that organic matter concentrates derived from humic ecosystems may consist of a diverse and temporally variable mix of carbon compounds leading to unreproducible results. Therefore, large-scale browning experiments (mesocosm or whole-ecosystem experiments) tend to rely on commercially available products as experimental browning agents. Most commonly, leonardite (i.e., oxidized lignite) products are used, which were originally manufactured for agricultural applications such as soil management or feed amendment (Quilty and Cattle 2011). In experiments, these products have been assumed to mimic the natural browning phenomenon, by being fairly recalcitrant and of poor nutritional quality while having similar physical and chemical properties as those of indigenous terrestrial DOM (Lennon, Hamilton, Muscarella and others 2013), or by being considered inert browning agents with no significant impact on the total bioreactive carbon (Lebret, Langenheder, Colinas and others 2018). However, there are indications that the use of these leonardite products may compromise the original purpose of their application in browning studies. For example, Urrutia-Cordero, Ekvall, Ratcovich and others (2017) reported the need to frequently re-supply the leonardite product HuminFeed during the course of an experiment in order to maintain the desired increase in water color. Lennon, Hamilton, Muscarella and others (2013) also described high flocculation rates of the leonardite product SuperHume when used in alkaline ponds as sinking of particles exported 5-12% of the total dissolved organic carbon (DOC) pool daily to the sediment.

Indeed, environmental conditions affect the behavior of browning agents in both natural and experimental settings. In lakes, formation of particles can be promoted by, for example, sunlight (Porcal, Dillon and Molot 2013; von Wachenfeldt, Sobek, Bastviken and others 2008), low pH, microbial activity (von Wachenfeldt, Bastviken and Tranvik 2009) and high concentrations of multivalent ions, in particular Ca^2+^ and Mg^2+^, which are typical for high alkalinity (i.e., hard water) lakes (Abate and Masini 2003). In addition, the fate of DOM compounds in freshwater ecosystems depends on their chemical composition, affecting their susceptibility to both photochemical and biological degradation (Kellerman, Dittmar, Kothawala and others 2014; Mostovaya, Hawkes, Köhler and others 2017). Sunlight mediated photoreactions can both completely mineralize DOC molecules or modify their bioavailability through the alteration of the molecular structure (Moran and Zepp 1997; Wetzel, Hatcher and Bianchi 1995). As the nature of the leonardite products used in browning experiments is largely unknown, the consequences of their exposure to sunlight and other environmental conditions are unpredictable and largely unknown.

The bioavailability of the browning agents used in manipulation studies and their effects on the microbial loop are crucial aspects that must be considered when evaluating their suitability in browning experiments. While the terrestrial DOM responsible for natural browning of freshwaters contains a labile fraction that serves as a carbon source for heterotrophic bacteria (Guillemette, McCallister and del Giorgio 2016), the leonardite browning agents used in experiments are believed to rather exclusively mimic the water color changes of the browning process (Lebret, Langenheder, Colinas and others 2018). However, while the effects of browning agents on bacterial communities and their functions remain unknown, Lennon, Hamilton, Muscarella and others (2013) showed that certain bacterial strains can use leonardite browning agents as a sole carbon source.

Additionally, leonardite products, such as HuminFeed and SuperHume, may contain compounds that are harmful or even toxic to organisms at higher trophic levels. Saebelfeld, Minguez, Griebel and others (2017) reported that HuminFeed negatively impacts reproduction and causes stress response in cultures of the cladocerans *Daphnia magna* and *Daphnia longispina*. In contrast, Lennon, Hamilton, Muscarella and others (2013) did not observe any negative effects of SuperHume on cultures of Daphnia *pulex* x *pulicaria*. Instead, they found a slight increase in fitness due to an earlier age at first reproduction.

While there are indications that leonardite browning agents interfere with bacteria and zooplankton, their impacts on phytoplankton have not been adequately studied. In the browning context, the invasive microalgae *Gonyostomum semen* (Raphidophyceae) is of particular interest as it causes extensive blooms in brown-water lakes (Rengefors, Weyhenmeyer and Bloch 2012). Thus, there is a strong ecological and societal interest in understanding the factors influencing the mass development of this algae by conducting experimental studies under browning conditions, potentially using leonardite products.

This study aims to assess the feasibility of the use of different browning agents commonly applied in aquatic browning manipulations by comparing their effect under different environmental conditions. This is the first time that the effects of two commercially available leonardite browning agents that are widely used in aquatic manipulation studies (i.e., “HuminFeed”, hereafter called HF, and “SuperHume”, hereafter called SH), are compared with a reverse osmosis concentrate extracted from a humic aquatic ecosystem (“Reverse Osmosis”, hereafter called RO). We characterized the browning agents chemically and tested whether they would act as an inert carbon source or if they would be bioavailable and, thus, subsidize the food web. Therefore, a mesocosm study was conducted to test responses at semi-natural scale, and several complementary laboratory experiments addressed specific processes. We assessed effects of the browning agents on both abiotic (chemical diversity and particle formation of organic matter), and biotic parameters (including bacterial production (BP), bacterial community composition (BCC), phytoplankton growth, as well as zooplankton abundance and life history) (see Table 1 for a summary of experiments conducted).

**Table 1:**
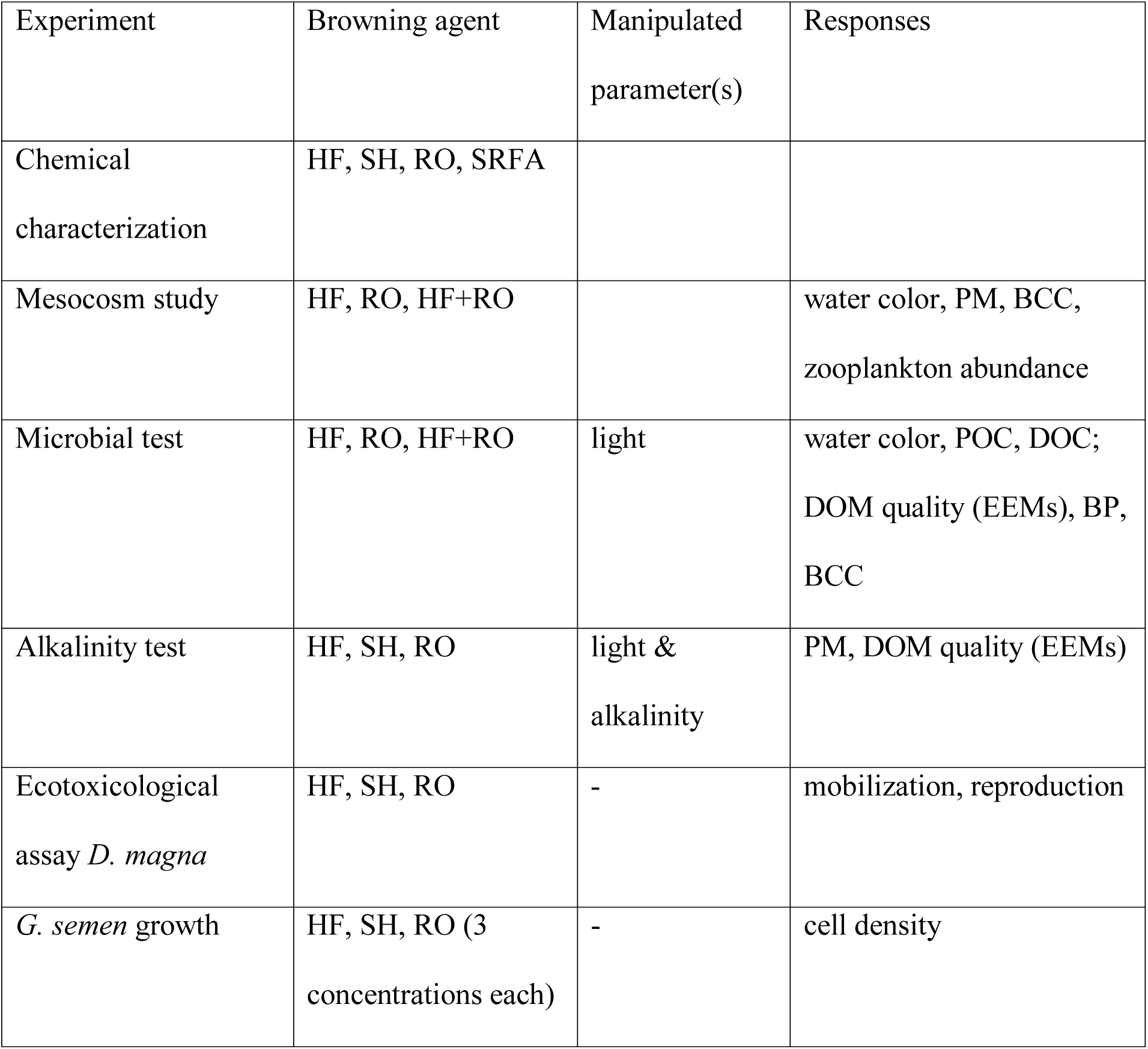
Overview of experiments for evaluating the effects of browning agents. HF= HuminFeed, SH= SuperHume, RO= Reverse osmosis concentrate, SRFA= Suwannee river fulvic acid, PM= particulate matter, BCC= bacterial community composition, EEM= Excitation emission matrix, BP= bacterial production.

## Materials and Procedures

### Description of browning agents

HuminFeed (HuminTech GmbH, Grevenbroich, Germany) is a commercially available food supplement for animal livestock. It is a water-soluble dry powder produced from alkaline extraction of oxidized lignite (leonardite). According to the manufacturer it consists of 82% humic substances, 18% compounds of lower molecular weights and no polysaccharides. To our knowledge, this agent has only been used in browning studies across Europe (e.g. Lebret, Langenheder, Colinas and others 2018; Meinelt, Paul, Phan and others 2007; Saebelfeld, Minguez, Griebel and others 2017; Urrutia-Cordero, Ekvall, Ratcovich and others 2017).

SuperHume (CropMaster, United Agricultural Services of America, Inc., Lake Panasoffkee, Florida, USA), another commercially available leonardite product, is a liquid containing 4% fulvic and 8% humic acids according to the manufacturer’s specification. This browning agent has been used in several studies of the browning phenomenon in Northern America (e.g. Lennon, Hamilton, Muscarella and others 2013; Muscarella, Jones and Lennon 2016; Weidel, Baglini, Jones and others 2017).

For comparison, we also used a reverse osmosis apparatus to produce a humic DOM concentrate from water collected from a local humic stream draining a forested wetland (59°55’0.5.0”N, 17°20’49.3”E). After an initial filtration through 0.2 μm pore size membrane filters and subsequent passage through a cation exchange resin (Dowex® 50W X8, Dow Chemical Company, Midland, MI, USA) the stream water was concentrated by reverse osmosis using a Real Soft PROS/2S unit (RealSoft, Norcross, GA, USA) as described by Serkiz and Perdue (1990), to a final concentration of approximately 800 mg C L^-1^. To obtain sufficient concentrate for our mesocosm experiment, we processed 3900 L of stream water that had a concentration 38 mg C L ^-1^, which required approximately 90 hours of on-site filtration.

### Chemical characterization of browning agents

We analyzed metals in digested samples of HF, SH and RO by inductively coupled plasma adsorption emission spectroscopy (ICP AES) using a Spectro Ciros CCD ICP-AES (Spectro, Kleve, Germany) as described in Appendix 1a. To identify the chemical properties of the different browning agents, we used Nuclear Magnetic Resonance (NMR) to determine proton chemical environments using a Bruker advanced Neo 600 MHz spectrometer (^1^H NMR: 600.18 MHz), equipped with a cryogenic tippled resonance probe TCI (CRPHe TR-1H &19F/13C/15N 5mm-EZ) as described in Appendix 1b. To measure the size, charge and mass distribution of the material High Pressure Size Exclusion Chromatography – High Resolution Mass Spectrometry (HPSEC-HRMS) was conducted with an Agilent 1100 HPLC (Agilent, Santa Clara, CA, USA) equipped with a UV-Vis Diode Array Detector for sample light attenuation (Agilent 1100, Santa Clara, CA, USA) and an Orbitrap mass spectrometer (LTQ-Velos Pro, Thermo Fisher Scientific, Waltham, MA, USA) in series that detected negatively ionizable molecules by electrospray ionization mass spectrometry, as described in Hawkes, Sjöberg, Bergquist and others (2019) and Appendix 1c. Solutions of HF, SH, and Suwannee River Fulvic Acid (SRFA, International Humic Substances Society, Batch 2S101F) in deuterated water (99.96%, Eurisotop) were prepared to 4.3, 5.6, and 1.25 mg ml ^-1^, respectively. We used SRFA instead of RO because it is available in powder form, facilitating dissolution in deuterated water. Due to the similar production process, we do not assume important differences between the two samples – both are constituted by typical aquatic DOM. More details of the chemical characterization methods can be found in the Supplementary material Appendix 1.

### Mesocosm study: in situ responses to browning agents

The effect of two different browning agents (HF and RO) and their combination (i.e., HF+RO) was assessed by a mesocosm experiment implemented for four weeks between June 15^th^ and July 13^th^ in 2016. The mesocosm facility consisted of 20 high-density polyethylene, white opaque, open top cylinders of 2 m depth and a diameter varying between 92 and 101 cm. It was located in Lake Erken (59°50’09.6’’N, 18°37’52.3’’E), held and fixed to a floating wooden jetty close to the lake shore. Details of the experimental set-up can be found in Nydahl and others (2019). In short, after filling the mesocosms with lake water, four treatments with five replicates of the following DOC concentrations (mean ± standard error) were established: Control (13.0 ± 0.05 mg C L ^-1^), HF (18.4 ± 0.06 mg C L ^-1^), RO (18.1 ± 0.10 mg C L ^-1^), and HF+RO (23.5 ± 0.05 mg C L ^-1^). Every week an integrated water sample of 15-18 L was collected from each mesocosm using a 1.5 m long tube sampler, to analyze water color, particle formation (i.e., particulate matter (PM) formation), BCC, and zooplankton abundance.

Zooplankton samples were collected by filtering 5 L of water through 55 µm plankton net, and preserving the zooplankton in Lugo’s solution. Zooplankton was counted and species abundances were determined using an inverted microscope (Leica, DM, IL LED Fluo, Leica Microsystems GmbH, Wetzlar, Germany). The immediate impact of the two browning agents on the abundance of Copepoda and Cladocera was evaluated at the first sampling campaign, i.e. approximately 16 hours after the addition of the browning agents.

### Microbial test: Effects on bacteria and interaction of light and browning agents

In order to assess the effect of light exposure on the browning agents, a laboratory scale experiment was performed with similar treatments as in the mesocosms but with different light conditions (hereafter called the microbial test). Four one-liter replicates of the HF, RO, HF+RO and Control treatments (initial DOC concentration: 18.0, 22.2, 27.1 and 12.9 mg C L^-1^, respectively) were placed either in ambient day light at a window facing west or in the dark for 22 days. Light and dark treatments were both performed at room temperature. Prior to the addition of the browning agents (HF and RO) the water was filtered through Whatman GF/F filters to remove larger particles, microeukaryotes, zooplankton, and phytoplankton. All treatments were sampled for bacterial production (BP) and water color at six time points (start, 6 h, 24 h, 120 h, 336 h and 528 h), and for particulate organic carbon (POC), dissolved organic carbon (DOC)), and DOM quality measurements by Fluorescence Excitation Emission Matrix (EEM) spectroscopy at the beginning and end (0 h and 528 h). BCC was assessed only at the end of the experiment (528 h).

### Alkalinity test: Interaction of water hardness and browning agents

To assess whether the interaction of light and browning agents depends on water hardness (measured as alkalinity and conductivity), and to compare the effects of the two most commonly used leonardite browning agents (i.e., HF and SH), a second laboratory experiment was performed (hereafter called the alkalinity test). The experiment was conducted using the three different browning agents (HF, SH, RO) added to water from Lake Erken, which is characterized by hard water (alkalinity: 1.81 meq L^-1^; conductivity: 27.4 mS m^-1^ -average of 25 years of monitoring), or to water from Lake Ljustjärn (59°55’23.1’’N, 15°27’18.5’’E), characterized by soft water (alkalinity: 0.08 meq L^-1^; conductivity: 4.45 mS m-1; (Sobek, Algesten, Bergstrom and others 2003). Prior to the experiment, the lake waters were prefiltered through a 50 µm mesh-size plankton net to remove zooplankton and larger particles. For the browning agent treatments (HF, SH, RO) 10 mg TOC L^-1^ of each of the agents were added to water from both lakes, respectively, and then incubated in light or dark. The light treatment was performed by incubating the bottles first outside in natural sunlight for 7 days (temperature between 8 and 22 °C), and subsequently in a dark constant temperature room (20 °C) for another 7-8 days. The dark incubations were kept for the entire experiment (14 days) in the dark constant temperature room. All treatments were performed in 500 mL glass bottles in triplicates. All bottles were sampled for PM concentration and DOM quality assessment by fluorescence EEM spectroscopy measurements on the first and last day of the incubations.

### Ecotoxicological assay: zooplankton life history responses

To assess the effect of the browning agents on the life history of zooplankton we used an acute immobilization test (OECD standard 202) and a reproduction test (OECD standard 211) with lab cultures of *D. magna*. The daphnids originated from a single clone (environmental pollution test strain *Klon 5* of the State Office for Nature, Environment, and Customer protection North-Rhine Westfalia, Bonn, Germany) and were cultured in glass beakers containing M7 media (OECD standards 202 and 211) under a constant temperature of 20°C, and a 16:8 hours light:dark cycle. The animals were fed three times a week with 0.1-0.2 mg C Daphnia ^-1^ day ^-1^ of the green algae *Pseudokirchneriella subcapitata*. These algae were cultured in culture medium (OECD standards 201) with air bubbled into the culture under constant daylight conditions and temperature (20°C). Algal concentrations were determined using a flow cytometer (Parctec CyFlow Space, Goerlitz, Germany).

The immobilization test was carried out for 48 hours under constant temperature and light cycle (as described above). No food was provided during this test. For each browning agent (diluted in M7 medium) and a control (pure M7 medium), four replicate vials were adjusted for browning agent concentration of 5, 10, 20, and 30 mg C L^-1^. Five individual neonates born within 24 h were placed in each vial containing 10 ml of the respective treatment solution. After 24 and 48 hours, the number of immobilized daphnids were recorded.

The reproduction experiment was carried out for 21 days under constant temperature, light cycle, and food conditions (as described above). The browning agents were amended to M7 medium to a concentration of 10 mg C L^-1^ and a control was set up with pure M7 medium. Twelve replicate vials were adjusted for a concentration of one neonate per vial. Each day the daphnids were removed from the vials, separated from their offspring (if applicable) and offspring were counted before returning the experimental daphnids to their respective vials. Medium or browning agent was refreshed four times during the period of the experiment.

Net reproduction rate (R0) was calculated over the 21 days of the experiment using the formula:

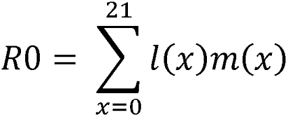

where *l*(*x*) is the number of individuals surviving to age x (in days), and *m*(*x*) is the number of offspring per surviving female between age x and x + 1. Furthermore, number of offspring per clutch, total number of offspring, age at first clutch, number of clutches and number of offspring at first clutch of *D. magna* was estimated.

### Growth dynamics of Gonyostomum semen

We tested the response of *G. semen*, a phytoplankton species known to be associated to high water color (Cronberg et al. 1988, Rengefors et al. 2012), to the different browning agents. The experiment was performed using a monoclonal strain of *G. semen* isolated from the humic lake, Koreivienė and others 2016). The strain was grown in batch mode with an initial cell density of 250 cells ml^-1^ and a total volume of 30 ml per cell culture flask (Thermo Scientific Nunc, Rochester NY, United States) under constant light intensity (100 μmol photons m^-2^ s^-1^ in a 14:10 hours light:dark cycle) and constant temperature of 20°C. Three different concentrations of the three browning agents (HF, SH and RO) were used to test their effect on *G. semen* growth rates compared to a control (MWC+Se – Wright’s cryptophyte medium MWC modified from Guillard and Lorenzen (1972), and with an addition of 4.5 nM Na_2_SeO_3_. Concentrations of the three browning agents, dissolved in MWC+Se medium, were set to low (2.4 mg l^-1^), medium (7.2 mg l^-1^) and high (21.6 mg l^-1^) levels of DOC. Each treatment including the control had five replicates. Cell density was determined after 12 days using a FlowCam Benchtop B3 (Fluid Imaging Technologies Inc., Scarborough ME, United States) equipped with a 300 μm flow cell, and specific growth rates *μ* per day during the exponential growth phase were calculated from the obtained cell densities as 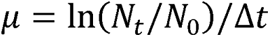.

### Chemical analyses of experiments

Prior to water color, DOC and EEMs analyses, the water samples were filtered through pre-combusted GF/F filters (Whatman, GE Healthcare, UK). Water color was measured as absorbance at 440 nm and 420 nm using a Lambda 40 UV-visible spectrophotometer (Perkin Elmer, Waltham, MA, United States). DOC concentration was measured on a Total Carbon Analyzer (Sievers M9 Laboratory Analyzer, GE Analytical Instruments, Boulder, Colorado, USA), while EEM spectroscopy for qualitative assessment of DOM was performed as described before (Kothawala, Stedmon, Müller and others 2014). Briefly, the UV-visible absorbance spectra were determined using a Lambda 40 UV-visible spectrophotometer (Perkin Elmer), while EEMs were obtained using a fluorescence spectrophotometer (SPEX FluoroMax-4, Horiba Jobin Yvon, Kyoto, Japan). Milli-Q water was used as blank and its values were subtracted from the EEM, which were also corrected for instrument biases and inner filter effects.

Samples for particulate matter analyses were collected on pre-combusted glass microfiber filters. For total PM quantification the weight of the empty filters was extracted from the weight of dried filters. For particulate organic carbon (POC) analysis in the light test, the samples were collected on GF/F filters and acidified with 10% HCl after filtration to remove inorganic carbon prior drying in an exicator. Subsequently, POC was analyzed using an Elemental Combustion System (Costech Instruments, Cernusco s/Nav., Italy). Total organic carbon (TOC) was calculated as the sum of POC and DOC.

### Microbial analyses of experiments

The BCC was assessed by filtering approximately 250 ml of water onto 47 mm diameter 0.2 μ pore-size polyether sulfone (PES) membrane filters (Supor-200, Pall Corporation, Port Washington, NY, USA). DNA was extracted from the filters and amplified, sequenced, and the raw sequences were processed as in Segura, Nilsson, Schleucher and others (2019). Briefly, the V3-V4 region of the bacterial 16S rRNA gene was amplified and sequenced on an Illumina MiSeq platform at National Genomics Infrastructure (NGI, SciLifeLab, Uppsala, Sweden) and the raw sequences were processed into operational taxonomic units (OTUs) using the UNOISE pipeline (Edgar 2016). Samples with less than 5000 reads were removed, leaving 29 samples for the final data analyses. Prior to these analyses, all remaining samples were rarefied to the sample with lowest read count.

Heterotrophic bacterial production (BP) was determined immediately after sampling via the measurement of the incorporation rate of L-^3^H-leucine (Perkin Elmer, Waltham, Massachusetts, USA, specific activity 161 Ci mmol^-1^) into the protein fraction based on the protocol of Smith and Azam (1992) as in Székely, Berga and Langenheder (2013).

### Data analyses

Processing of the EEMs was performed using MatLab (MatLab 7.7.0, The MathWorks, Natick, USA) and the *FDOMcorr* toolbox (Murphy, Butler, Spencer and others 2010) as described before in Kothawala, Stedmon, Müller and others (2014). Based on Fellman, Hood and Spencer (2010), the specific peaks C, A, T, B and M were extracted for further qualitative analyses of DOM. All statistical analyses were performed using the R (version 3.4.3) environment for statistical computing (R Core Team 2018). The effect of the different treatments was assessed by comparing either the parameters measured at the end of the experiments or the changes of the parameters during the experiments by calculating the difference between the final and initial values. The importance of the different treatments was estimated by linear models tested by analyses of variances (ANOVA). Alternatively, to test the effect of the treatments in time for PM and for BP in the case of the mesocosms experiment and the microbial test, respectively, a mixed effect model repeated measures ANOVA was performed using treatment, time and their interaction as fixed effects and mesocosm ID as random factor. Significant differences between treatments were determined by coefficients of the model or Tukey’s post hoc analyses. To fulfill the assumptions of the applied ANOVAs data was log-transformed when necessary (number of offspring at first clutch of *D. magna*, POC in the microbial test, PM in soft water of the alkalinity test), or inverse-transformed (age at first clutch of *D. magna*).

Ordination of multivariate data was implemented by non-metric multidimensional scaling (NMDS) with Euclidean and Bray-Curtis dissimilarity indexes for the change in the specific peaks of EEMs (i.e., A, B, C, M, T peaks) and bacterial OTUs, respectively. The importance of the different treatments was assessed by permutational analyses of variances (PERMANOVA, 999 permutations). All multivariate analyses were performed using the vegan package of R version 3.6.1. (Oksanen, Blanchet, Friendly and others 2017).

## Assessment

### Chemical characterization of browning agents

Compared to SH and RO, HF had elevated levels of aluminum, iron and sodium, while SH had higher concentrations of calcium compared to HF and RO, and higher aluminum and iron than the RO (Table 2).

**Table 2:** Chemical composition of HuminFeed, SuperHume and Reverse Osmosis dry extracts in µg (mg C added)^-1^. LOD = Limit of detection.

| Browning agent | Al | As | Ca | Cu | Fe | Na | Ni | P | Zn |
| --- | --- | --- | --- | --- | --- | --- | --- | --- | --- |
| HuminFeed | 79.31 | 0.078 | 19.9 | 0.061 | 24.016 | 41.576 | 0.097 | 0.265 | 0.045 |
| SuperHume | 10.81 | <LOD | 84.9 | <LOD | 8.504 | 11.389 | <LOD | 0.021 | <LOD |
| Reverse Osmosis | 0.79 | <LOD | 45.3 | <LOD | 1.896 | 8.689 | <LOD | <LOD | <LOD |

NMR showed that HF and SH are both characterized by high abundance of aromatic protons in comparison to SRFA. The abundance of aliphatic ‘terpenoid-like’ protons (0-1.6 ppm) was similar for all three, and SRFA had the highest abundance of carboxylic rich alicyclic material and carbohydrates (Supplementary figure Appendix 2).

HPSEC-HRMS indicated strong light absorbance properties over the chromatographic separation of all three browning agents (Supplementary figure Appendix 3). Only the SRFA sample contained ionizable material in the range 350-450 Da, where DOC is typically found to be at a maximum in mass spectrometric analyses of organic matter from aquatic environments. The elution time of this agent was typical for DOM using this method (Hawkes, Sjöberg, Bergquist and others 2019), between 9-12 minutes. This result indicates that HF and SH do not contain carboxylic acids, with mass 200-800 Da, which are typical for DOM from aquatic environments – and this corresponds well to the NMR data, as these mixtures also contained little carboxylic rich alicyclic material (Hertkorn, Benner, Frommberger and others 2006). Instead, they are constituted by higher molecular weight aromatic compounds, which may explain their lower solubility and tendency to coagulate.

### Abiotic effects: water color and particle formation

In the mesocosm study, as expected, the addition of all browning agents (i.e. HF, RO and HF+RO) increased water color compared to the control (repeated measures ANOVA F _3,75_ = 1532, *p* < 0.001) and the increase was the most substantial for the treatments containing HF (i.e., HF and HF+RO, Figure 1a, Appendix 4). However, the color darkening effect of the browning agents decreased with time (effect of sampling time: F _4,75_= 4.768, *p* = 0.002) and the most substantial changes were detected at the beginning of the experiment between the first and the second sampling (Figure 1a). PM concentrations also varied among treatments (repeated measures ANOVA: F _3,16_ = 73.1, *p* < 0.001) with the highest concentrations also measured in the HF treatments (Figure 1b). Furthermore, PM concentrations changed over time (repeated measures ANOVA: F _4,64_ = 5.1, *p* < 0.001) with HF treatments showing increasing PM concentration until the third sampling.

**Figure 1:**
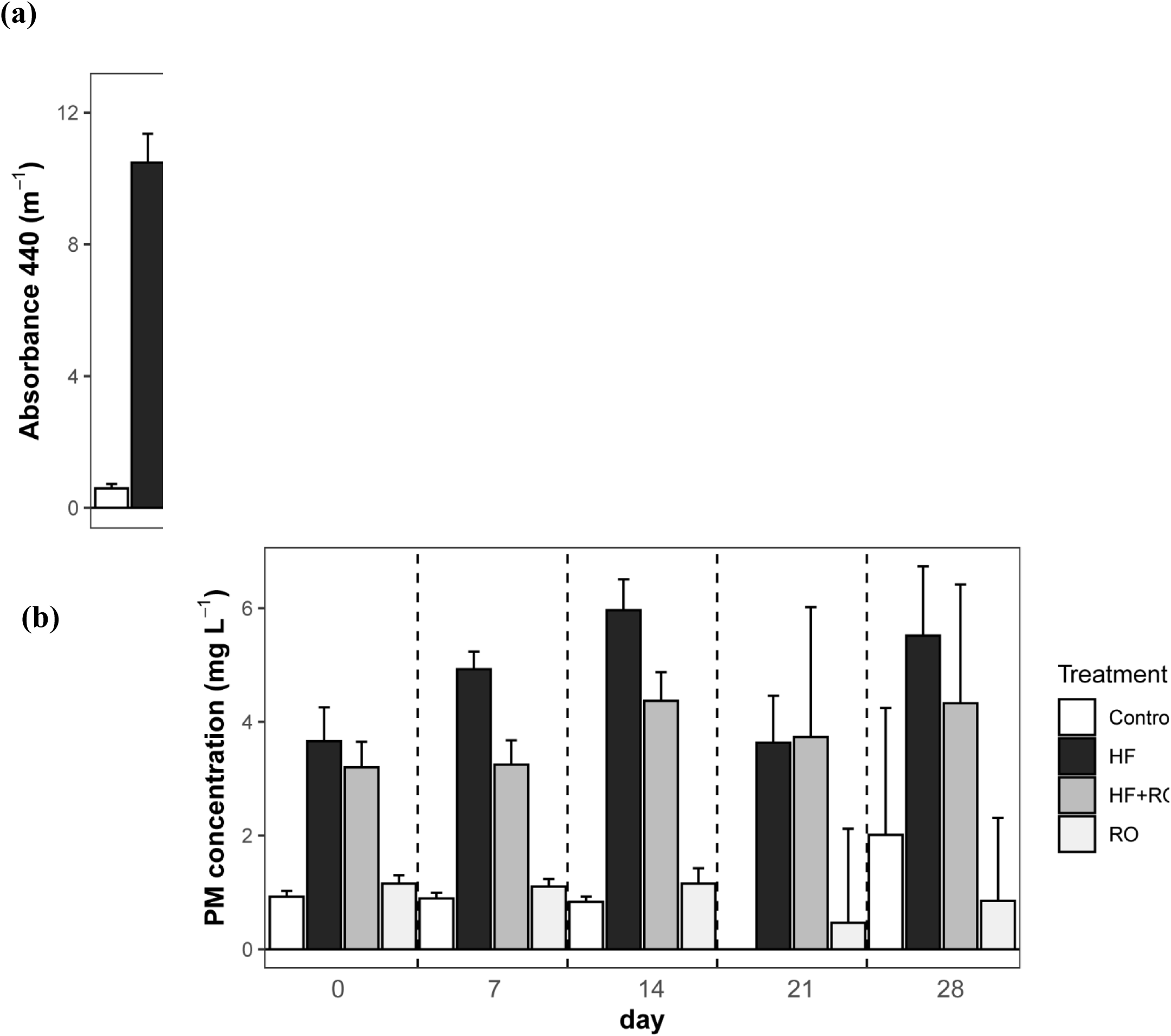
Changes and differences of a) water color (Abs_440_) and b) particulate matter (PM) concentrations of four different browning agent treatments (Control, HuminFeed (HF), Reverse Osmosis concentrate (RO), and the mix of the two (HF+RO)) during the four weeks of the mesocosm study.

In the microbial test using water from Lake Erken, the different carbon treatments (HF, RO, HF+RO and Control), the light treatment, and the interaction of the two all had a significant but variable effect on the change in water color, and POC and TOC concentration (Table 3). In the dark incubations, there was no decrease in the water color in any of the treatments (Figure 2, original absorbance values in Supplementary Figure Appendix 5 and statistical tests in Supplementary Material Appendix 6a/I), while in the light treatments, water color decreased in all treatments the decrease was significantly higher in the HF amended treatments (HF and HF+RO) than in the RO or control treatments (Figure 2a, Supplementary Material Appendix 6a/I). Regarding POC concentrations, in the dark, only the samples with added HF (HF and HF+RO) increased significantly in POC compared to the control (Figure 2b, Supplementary Material Appendix 6a/I), while, in the light POC significantly increased in all treatments compared to the dark controls with the highest increases in the samples with added RO (i.e., HF+RO and RO) (Figure 2b, Supplementary Material Appendix 6a/II). However, when DOC was considered or DOC and POC together as TOC, the picture was different. In the dark incubations, DOC loss was detected in all treatments (Figure 2c), while TOC loss was detected in both treatments with RO (HF+RO and RO), and in the control, but not in the treatment with only HF (Figure 2d). The most significant loss of both DOC and TOC was measured for the treatments with RO. However, in the light incubations, the detected DOC and TOC loss was opposite to the dark treatment with the highest losses seen in both of the HF treatments (i.e., HF and HF+RO) (Figure 2c, d). Thus, when the results of the different carbon analyses are combined, it is clear that while DOC decreased in all treatments, for most cases (except HF treatment in dark and control in light) this could not be explained solely by POC increase as TOC concentration also decreased (Figure 2b-d, Supplementary Material Appendix 8).

**Table 3:**
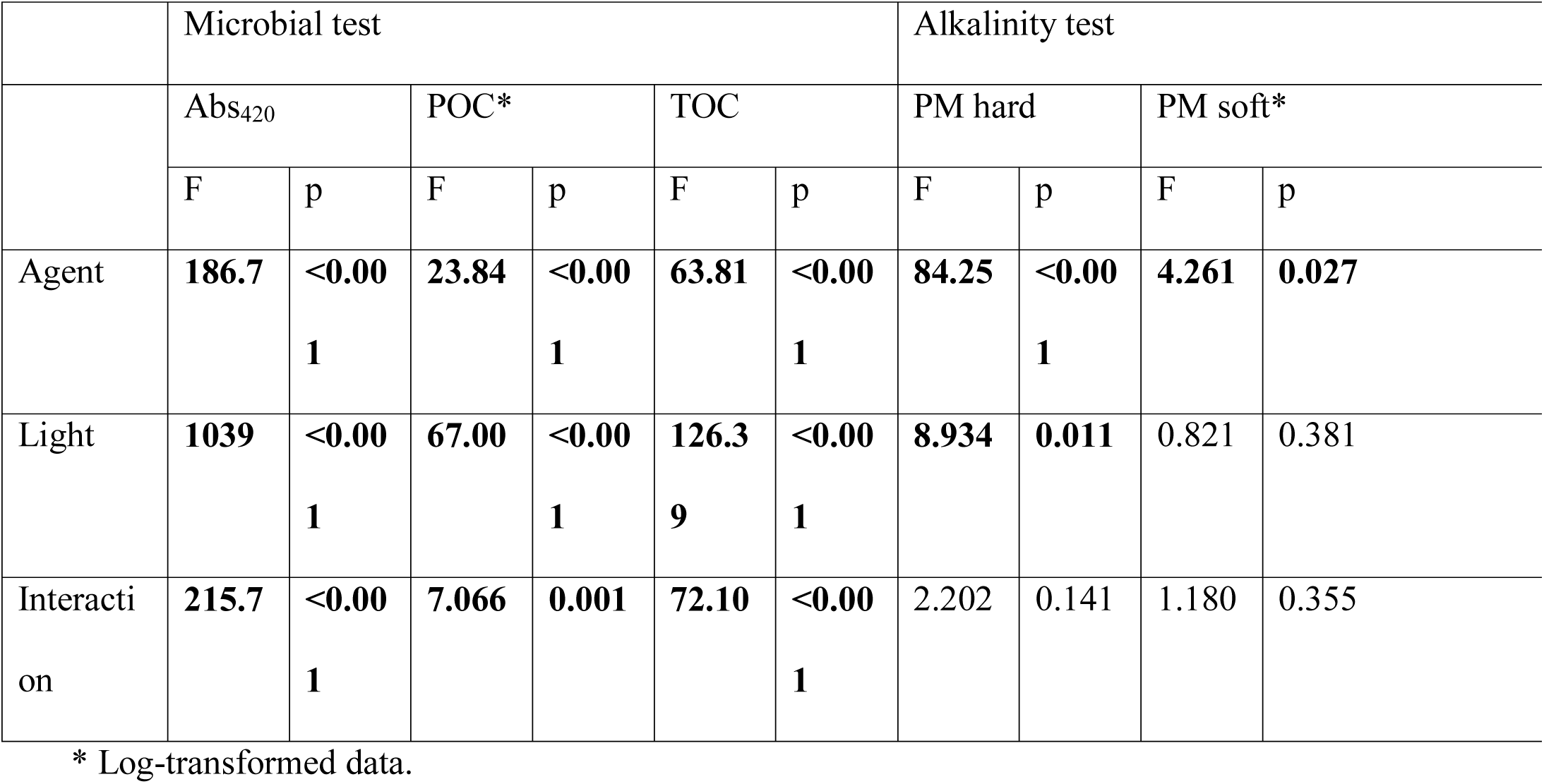
Water color changes (Absorbance at 420nm) and flocculation occurring in treatments of the different browning agents (HuminFeed, SuperHume and Reverse Osmosis). Results of ANOVAs from the microbial and alkalinity test. Bold font depicts significant effects with p < 0.05.

**Figure 2:**
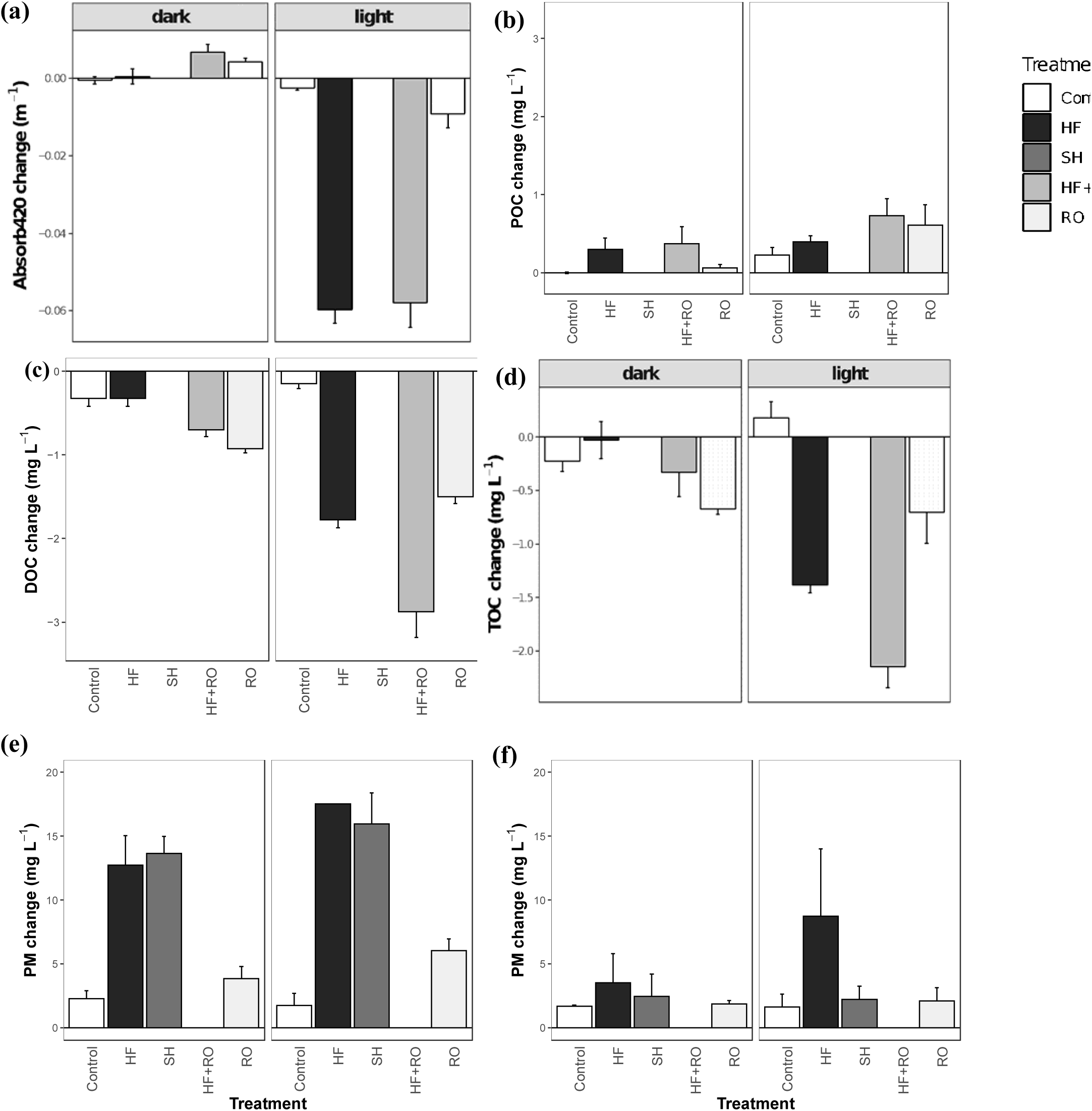
Changes in measured abiotic parameters throughout the microbial (22 days) and alkalinity test (14 days) under dark and light treatments using different browning agents (Control, Reverse Osmosis concentrate (RO), and leonardite containing treatments (HuminFeed (HF), SuperHume (SH), the mix of HF and RO (HF+RO)). For the microbial test, a) changes in water color (Abs_420_), b) particulate organic carbon (POC), c) dissolved organic carbon (DOC), and d) total organic carbon (TOC); and for the alkalinity test, changes in particulate matter (PM) in e) hard water and f) soft water are shown.

In the case of the hard water incubations (i.e., water from Lake Erken), PM was significantly affected by both the different added agents (i.e. HF, SH and RO) and the light conditions but not the interaction of the two types of treatments (Table 3). In both dark and light incubations, the highest increase of PM was measured for treatments with leonardite products (i.e. HF and SH). Although the PM increase was significantly higher in light than in dark, the difference between the dark and light treatment of the same agent was not significant (Figure 2d, Supplementary Material Appendix 6b/I). Unfortunately, the number of replicates decreased from 48 to 41 in the alkalinity test due to bottles breaking during the light incubations. In the case of soft water (i.e. water from Lake Ljustjärn) only the browning agents had a significant effect on the changes of PM, but not light treatment or the interaction of the two treatments (Table 3). The impact of browning agents was primarily driven by an outlier value in the light HF treatment (Figure 2e). However, no significant differences between pairwise comparisons could be detected (Supplementary Material Appendix 6b/II).

### Abiotic effects: qualitative DOM changes based on EEMs

The PERMANOVA of the change of extracted EEM spectroscopy peaks during the microbial and alkalinity tests revealed significant differences between the samples depending on the browning agents (levels: control, RO and leonardite containing treatments (i.e, HF, HF+RO, SH)), water hardness and the interaction of these two factors with light (Supplementary Material Appendix 9). The largest difference between treatments was detected for the peak related to substances with high molecular weight and aromatic humic nature (Peak A), in which all the treatments including leonardite (HF+RO, HF and SH) were distinct from the control in the light treatment in hard water (Supplementary Material Appendix 10a). Also the peak related to biological activity (Peak M) had the same trend with higher values detected for the treatments with leonardite in the light treatment in hard water, though the difference was less pronounced than for the peak A (Supplementary Material Appendix 10b). This was supported by the NMDS plot (Figure 3a), which also showed clear differences among the treatments as the samples in the light treatments diverged from the samples in the dark treatments and the direction of the divergence depended on the browning agent with leonardite treatments associated to divergence along the first axis of the NMDS and RO treatments diverging along the second axis of the plot. Furthermore, the divergence along the leonardite-associated axis depended on the hardness of the water with soft water treatments showing no substantial divergence from the corresponding dark incubation treatments. Finally, a lesser divergence appeared for the hard water leonardite treatments of the alkalinity test, where the light exposure was shorter than in the light test (seven compared to 22 days).

**Figure 3:**
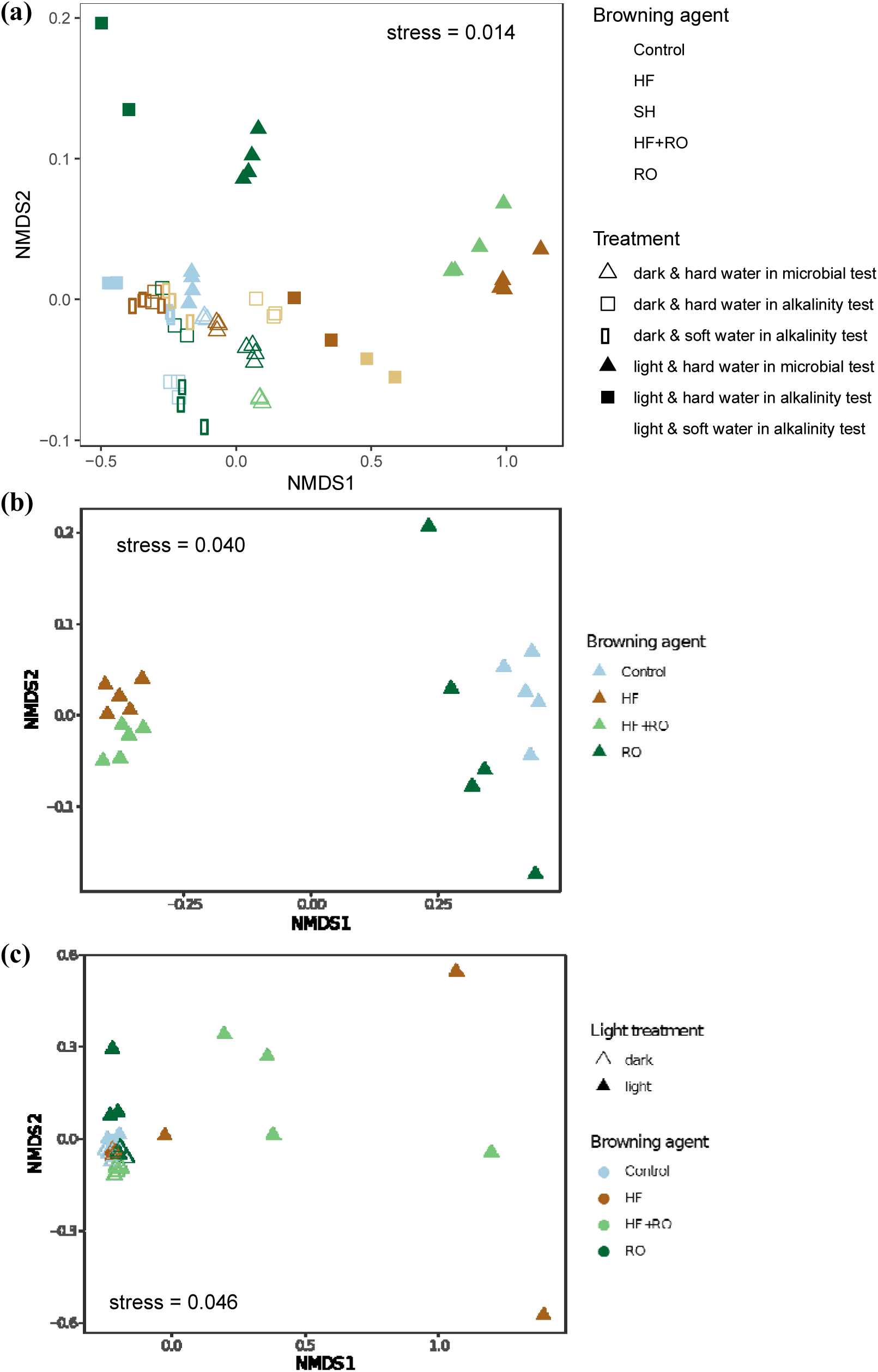
Ordination by non-metric multidimensional scaling (NMDS) of multivariate data. a) Ordination of the changes of the main peaks of the Excitation Emission Matrix (EEM) during the microbial and alkalinity test. b) Ordination of the bacterial community composition of the replicates at the end of the mesocosms experiment, and c) at the end of the microbial test. Blue color represents control, brown represents leonardite browning agents (dark brown HuminFeed (HF), light brown SuperHume (SH)), green represents treatments containing Reverse Osmosis concentrate (RO) (light green HF+RO, dark green RO). Empty symbols represent continuous dark treatment, filled symbols represent light exposed treatments. Hard water replicates are represented by triangle and square shape points, while soft water samples are symbolized by round symbols.

### Biotic effects: bacterial production and community composition

The PERMANOVA tests assessing the final bacterial community structure of both the mesocosms and the replicates of the microbial test revealed significant differences depending on the applied browning agent (Appendix 9). In addition, in the microbial test light and the interaction of light and browning agents was also significant. These results were supported by the NMDS plots of the two experiments (Figure 3). Specifically, by the end of the mesocosms study, the primary difference between the bacterial communities depended on the addition of the leonardite product HF, with HF and HF+RO mesocosms being clearly separated from the Control and RO mesocosms along the first NMDS axis (Figure 3b). Meanwhile, in the case of the microbial test, the interaction of light and the added browning agent was also reflected in the final bacterial community structure (Figure 3c) and their taxonomic composition of the browning agent treatments in light (Appendix 11). The communities that were incubated in dark did not differ substantially from each other, while for light incubations the Control treatments remained very similar to the dark incubations and RO treatments showed also only some minor differences in taxonomic composition and along the second axis of the NMDS, while the leonardite amended communities (i.e., HF and HF+RO) in the light treatment were substantially different from the other treatments and also presented greater variation among replicates (Figure 3c).

The heterotrophic bacterial production (BP) measured during the light test also displayed different trends for the dark and light incubations (Figure 4). The repeated measures ANOVA revealed significant effects of the different browning agent treatments (HF, RO and HF+RO) for both, the dark and light incubations (dark: F _3,12_ = 54.21,*p* < 0.001; light: F _3,12_ = 48.02, *p* < 0.001). However, in the case of the dark incubation there were no significant time-related differences (F _1,76_ = 0.659, *p* = 0.419; F _3,76_ = 0.397, *p* = 0.755), while incubation in light resulted in time dependent differences (F _1,76_ = 16.71, *p* = 0.001; F _3,76_ = 2.588, *p* = 0.059). More precisely, at the beginning of both dark and light incubations and throughout the dark treatment, the lowest BP values were measured for the HF treatments and the highest for the RO followed by the HF+RO (Supplementary Material Appendix 12a). In the light, however, BP in the HF treatment continuously increased with time and exceeded the control values already on the fifth day (120 h). In addition, the HF+RO incubations also showed different trends in light than in dark as they did not follow the declining trends of the RO incubations but instead became significantly higher by the end of the experiment (Supplementary Material Appendix 12b).

**Figure 4:**
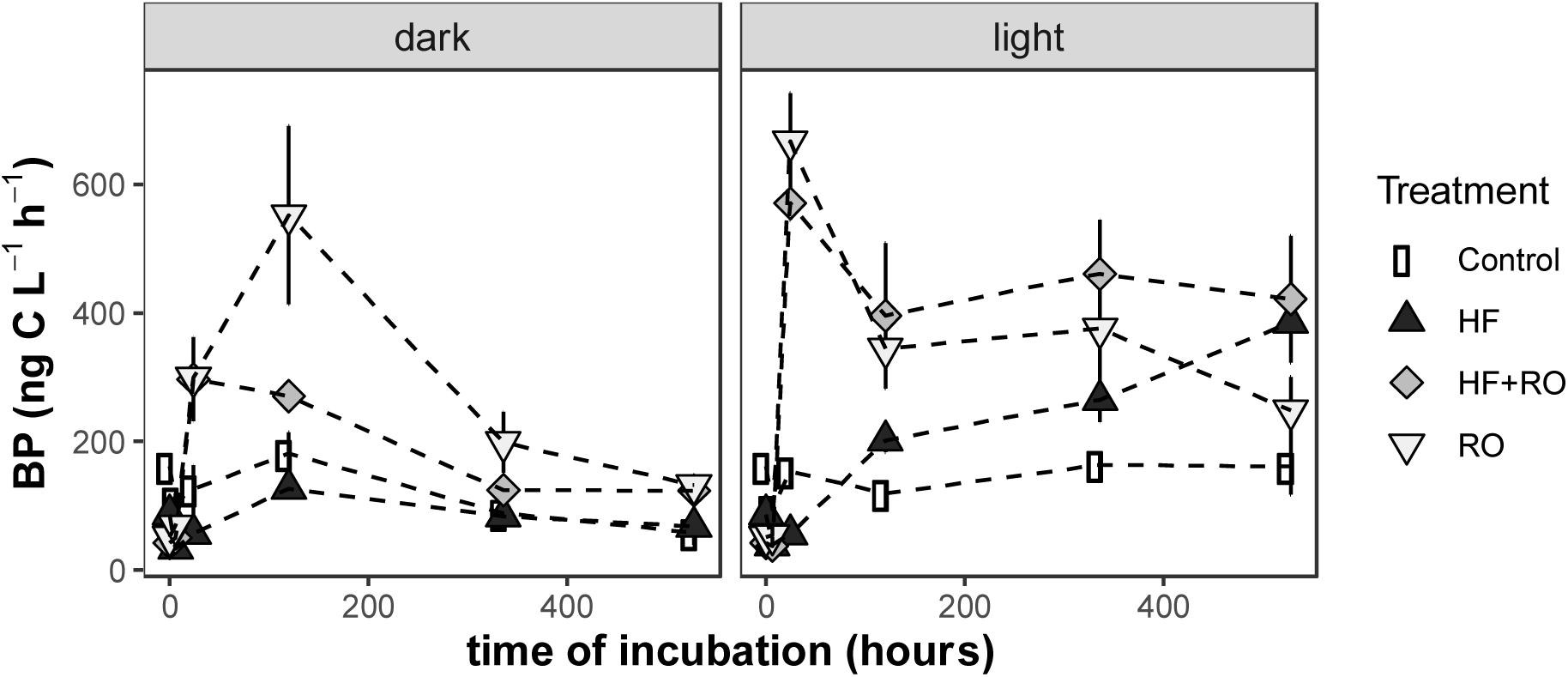
Changes in heterotrophic bacterial production measured by ^3^H-leucine incorporation during the microbial test for samples incubated in dark and light with different browning agents (Control, HuminFeed (HF), Reverse Osmosis concentrate (RO), and the mix of the two (HF+RO)).

### Biotic effects: zooplankton life history responses

The abundance of zooplankton was lower in mesocosms with HF (HF and HF+RO) compared to RO and Control (Figure 5), with overall significant treatment effects on the abundance of Cladocera (ANOVA: F _3,16_ = 4.4043, *p* = 0.026) and Copepoda (ANOVA: F _3,16_ = 11.86, *p* < 0.001). Coefficients of linear model depicted a significant lower abundance of Cladocera, and Copepoda in HF treatments (HF and HF+RO) compared to Control treatment, but the zooplankton abundances in the RO treatment were not significantly different to Control (Supplementary Material Appendix 13a).

**Figure 5:**
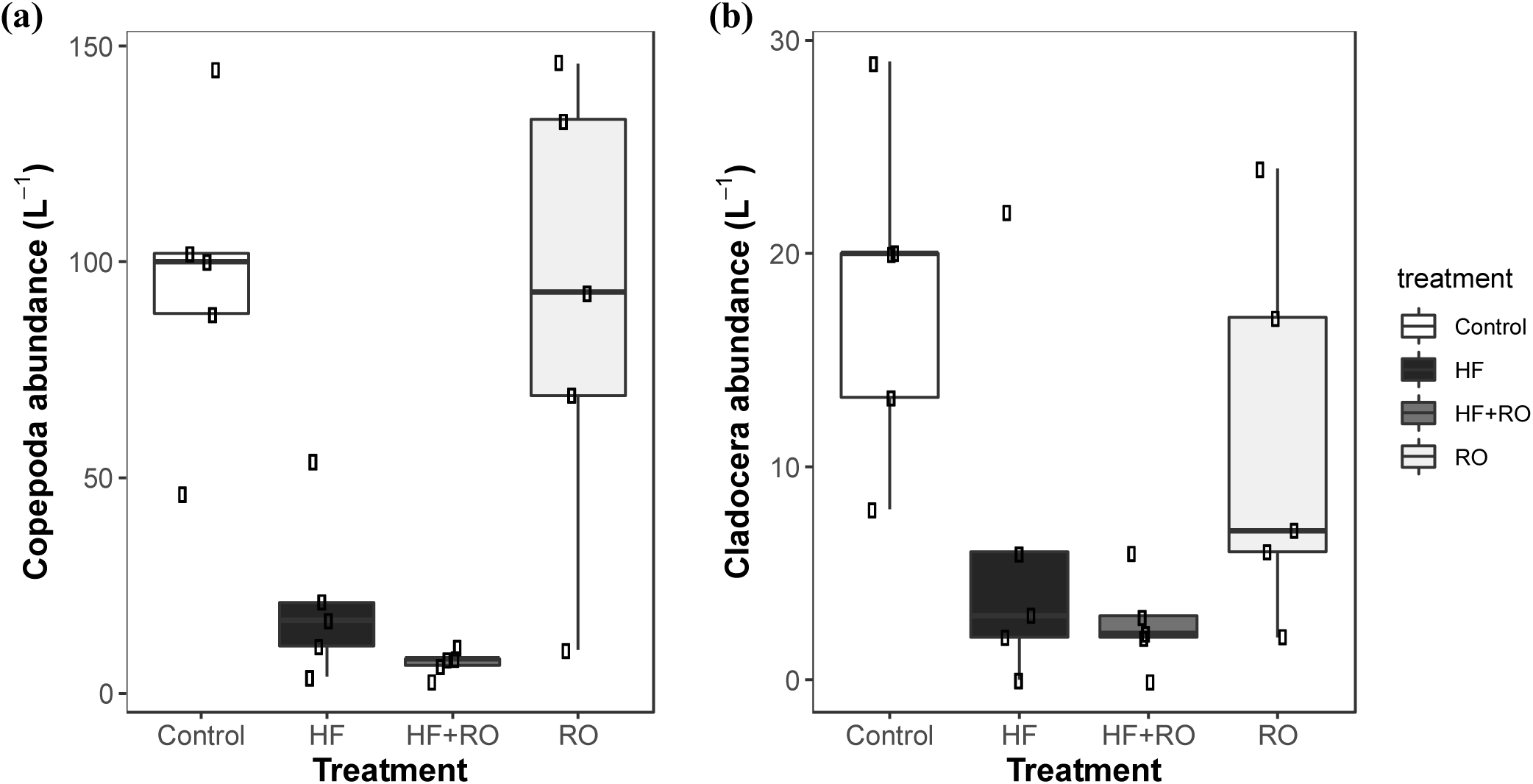
Mean abundance (± standard error) of a) Cladocera, and b) Copepoda in the mesocosms with different browning agent treatments (Control, HuminFeed (HF), Reverse Osmosis concentrate (RO), and the mix of the two (HF+RO)), approximately 16 hours after addition of the browning agents.

In the immobilization test, we could not observe an acute immobilization of *D. magna* during the 48-hour test period in any of the treatments. However, the different browning agents affected the reproduction of *D. magna* over the course of 21 days. Net reproductive rate (R0) that integrates both survival and fecundity was highest in the Control (64.1) and RO (57.7), and lowest in SH (52.5) and HF (45.9) treatments. The average number of offspring per clutch differed significantly between the treatments (ANOVA: *F*_3,25_ = 8.786, *p* < 0.001), with significantly lower numbers in all three browning agent treatments compared to the Control (Figure 6a, Supplementary Material Appendix 13b). The number of total offspring differed significantly between the treatments (ANOVA: *F*_3,25_ = 4.149, *p* = 0.002), with significantly lower number of offspring in the HF treatment compared to the Control (Figure 6b, Supplementary Material Appendix 13c). No significant differences were found between the treatments for age at first clutch, number of clutches, or number of offspring at first clutch.

**Figure 6:**
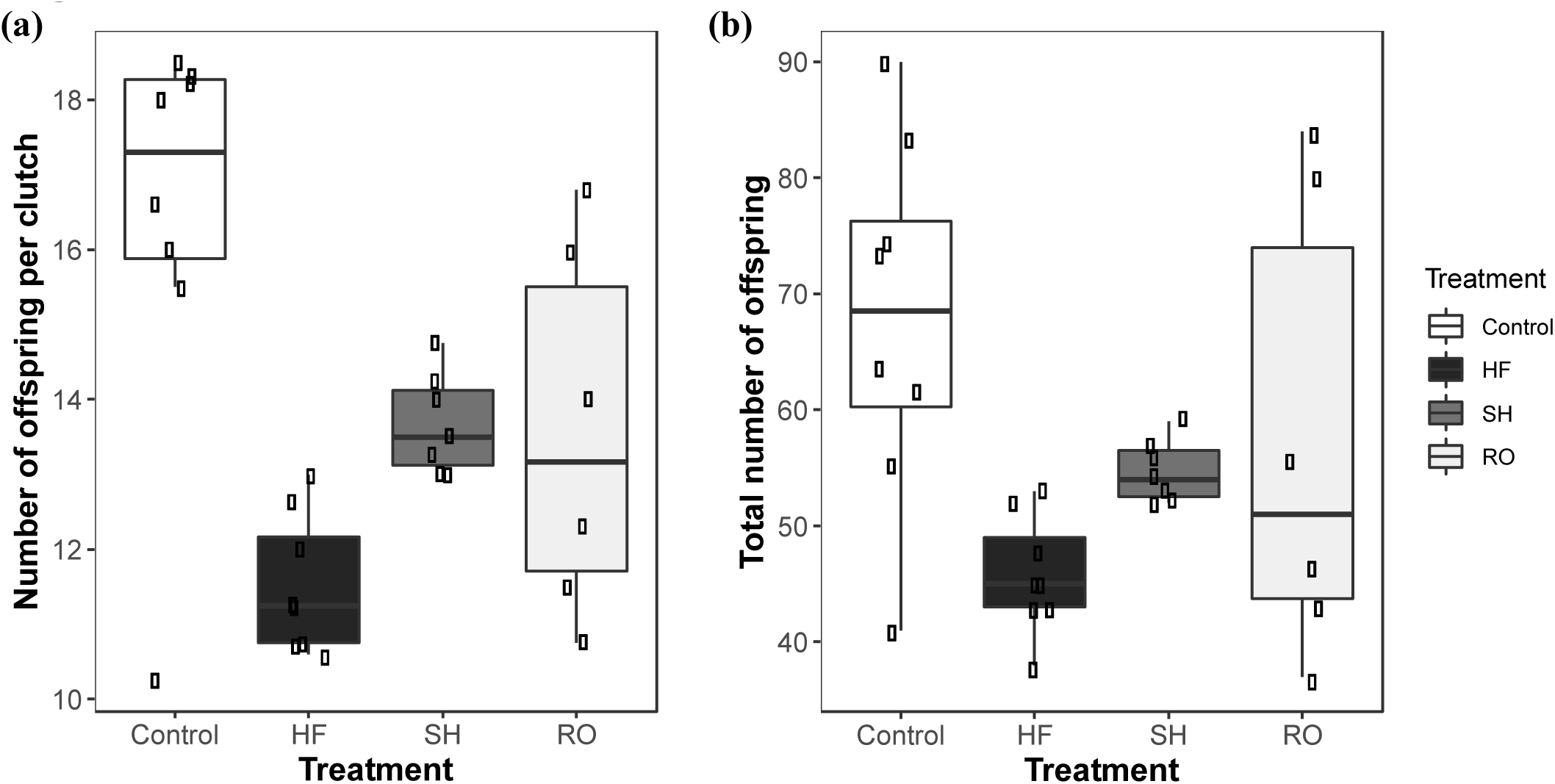
Fitness of *D. magna* after a 21-day reproduction test cultured under different browning agent treatments (Control, leonardite containing HuminFeed (HF) and SuperHume (SH), and Reverse Osmosis concentrate (RO)). a) Number of offspring per clutch, b) total number of offspring during 21 days.

### Biotic effects: growth dynamics of G. semen

Unfortunately, replicates with high RO concentrations were contaminated by coccoid green algae likely originating from the humic stream and could not be included in the analysis. Browning agents had a significant effect on growth rate of *G. semen* after 12 days (ANOVA: F _8,36_: 8.085, p < 0.001; Figure 7) and growth rate in replicates with medium concentrations of RO was significantly higher compared to the control, as indicated by the post hoc comparisons (Supplementary Material Appendix 14). Furthermore, the treatments with high concentrations of HF showed a significantly lower growth rate compared to the treatment with lower concentration.

**Figure 7:**
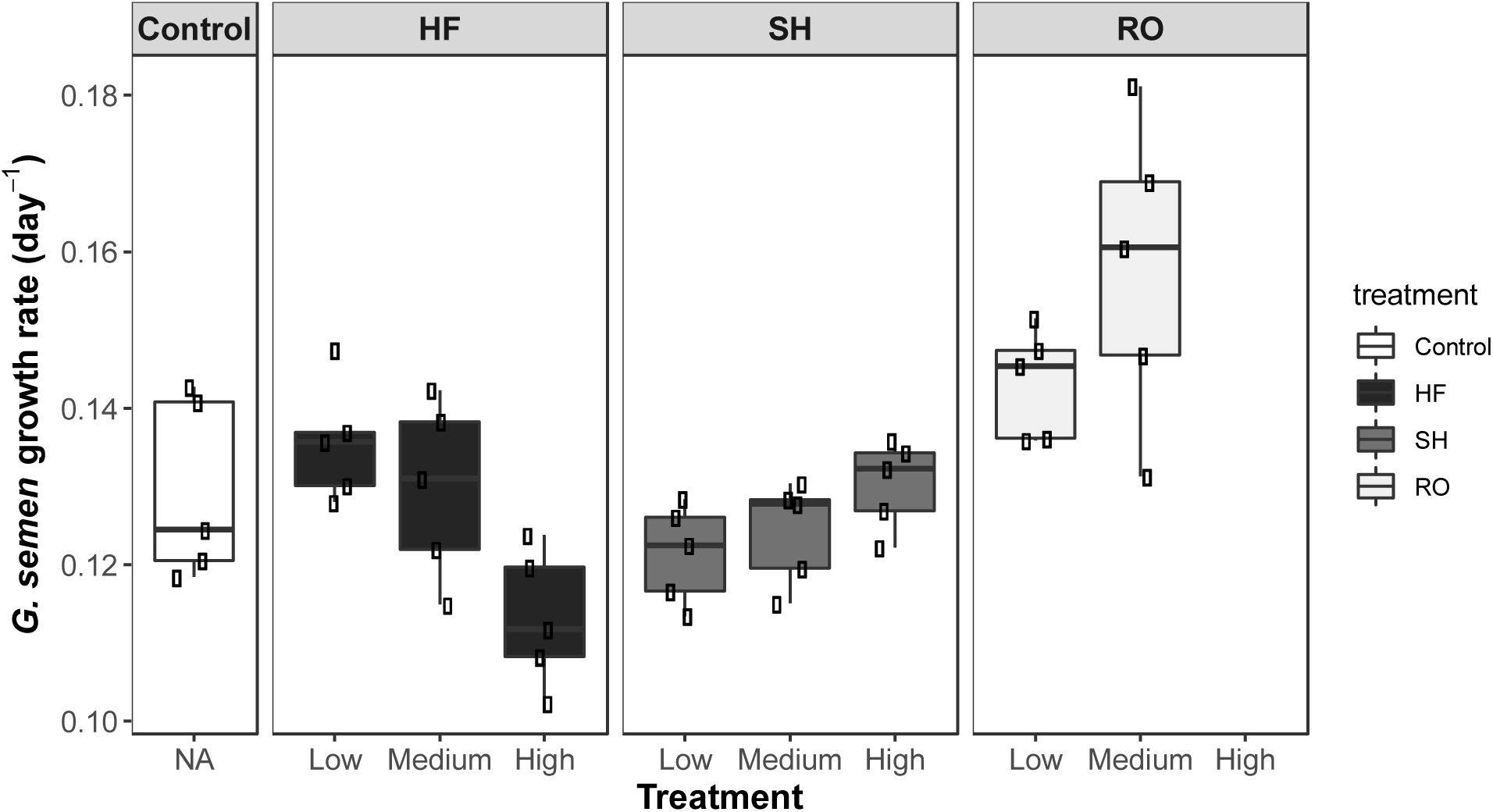
Growth rate of *Gonyostomum semen* under different treatments of browning agents (Control, leonardite containing HuminFeed (HF) and SuperHume (SH), and Reverse Osmosis concentrate (RO)) applied at different concentrations (Low, Medium, High) after 12 days. Exclusion of high concentrations RO treatments due to flagellate contamination.

## Discussion

In this study, we evaluated the suitability of leonardite products and a DOM concentrate obtained from a local aquatic environment in experimental studies of browning of freshwater ecosystems. We found that while leonardite products are very effective in establishing a light environment that mimics browning of surface waters, they have chemical characteristics that deviate from those of indigenous DOM. Consequently, leonardite agents may have biotic and abiotic effects that may bias conclusions on how browning affects ecosystems. Meanwhile, the tested DOM concentrate obtained from a local source via reverse osmosis, was less efficient when it came to water color changes but had less adverse effects on water quality and biota

### Chemical characteristics of browning agents

Both HF and SH had higher concentrations of aluminum, iron, sodium and phosphorus than RO. The multivalent cations, Al^3+^, Ca^2+^, and Mg^2+^, can stimulate particle formation and are applied as agents in drinking water purification (Matilainen, Lindqvist and Tuhkanen 2005). Although HF and SH are both leonardite products, they also had some differences with respect to the measured inorganic constituents. HF showed higher concentrations of all analyzed ions, but calcium was comparable to SH and RO. Of these, one of the most substantial differences was in aluminum, which was eight times higher in HF than in SH. Aluminum may be toxic for microbes and other organisms (Piña and Cervantes 1996), where toxicity increases with decreasing pH. In our experiments, both lake waters had neutral to alkaline pH, thus, toxicity of aluminum was limited. Further, aluminum toxicity is connected to iron availability with low iron concentration leading to more severe toxic effects of aluminum. In HF, high aluminum was combined with high iron, and that together with high pH suggests low potential for toxic impacts. Finally, HF also had higher phosphorus and iron concentration than SH, which, if it is bioavailable, could also stimulate microbial and phytoplankton growth.

The possible reasons for the variation in the composition between HF and SH could be that the leonardite used for their production comes from very different parts of the world (Germany vs. USA, respectively), and the fact that those are provided to the user in different forms. Therefore, it should be noted that especially when HF is used in manipulation studies, a range of compounds and elements are added that could provoke potential abiotic and biotic interactions.

Characterization of organic constituents of browning agents by NMR and HPSEC-HRMS identified HF and SH as highly distinct from material typically observed in humic waters (e.g. SRFA from the Suwannee River), as they both lack freely dissolved carboxylic acids that are typical components of naturally occurring DOC.

### Abiotic effects and their consequences

Both the mesocosm study and the microbial test demonstrated that the addition of both HF and RO increases water color compared to controls, with the browning effects of HF being much stronger than the effects of RO. For HF, the browning effect decreased in our mesocosms in accordance with a previous experiment, in which a weekly restocking of HF was needed in order to maintain a constant water color (Urrutia-Cordero, Ekvall, Ratcovich and others 2017). We also detected a water color decrease over time in the microbial test, but in this experiment, this only happened for the samples incubated in light, suggesting that photochemical reactions caused the color loss. Notably, the effect of light was detected in borosilicate glass bottles behind conventional window glass, where light levels were moderate and light at wavelengths <400 nm is very limited. Hence, most of the light that is expected to induce photochemical reactions in natural DOC was absent (Koehler, Broman and Tranvik 2016). As typical photosynthetic primary producers were absent from the microbial communities, biological photoreactions were also unlikely.

Besides photochemical reactions, particle formation of the browning agents and subsequent export through sedimentation could also have played a role in the loss of color, as we detected an increase in PM in the mesocosms and in the alkalinity test as well as POC increase in the microbial test. The increase in PM and POC was detected in both in dark and light and in almost all treatments including most of the controls, however, the increase was always the highest in treatments with leonardite agents. These results are in line with the findings of Lennon, Hamilton, Muscarella and others (2013) who estimated a loss of 5-12% of the total mass of SH due to particle formation. The alkalinity test further suggested an interaction between the browning agents and the different ion concentrations (i.e., alkalinity) of the lake waters, as both HF and SH caused more PM formation in the hard Erken water than in the soft Ljustjärn water (Figure 2 d, e). In the microbial test the significant effect of light on POC change was caused by higher POC formation in the control and RO samples, while the extent of POC was similar in light and dark for the HF and HF+RO treatments. Meanwhile in the alkalinity test the effect of light in particle formation was not significant for treatment pairs (e.g., SH in dark vs SH in light). Overall, all these suggest that particle formation in the leonardite treatments was not affected by light making it unlikely that water color loss happened due to this process.

The microbial test provided a more likely explanation for the substantial loss in color detected for H. TOC and DOC changes in the dark incubations showed that while all samples did decrease in DOC, this could be explained by flocculation only in the case of HF, while the control and the RO containing samples (i.e. RO and HF+RO) did experience organic carbon loss presumably due to mineralization of organic carbon through biodegradation. The fact that TOC loss in the HF treatments could be completely explained by POC formation and did not even reach the level of the controls suggests that HF was not only inert as suggested by some users (Lebret, Langenheder, Colinas and others 2018), but may have been inhibitory to the biological processes that could have degraded the background organic carbon in the water used for the experiment. This idea of inhibitory effects of HF was further supported by the effects on bacterial production. BP was low in the dark treatment of the microbial experiment, as the BP values of the HF samples remained below or close to the values of the control samples and also the values of the HF+RO were mostly below the samples containing RO agent only. Meanwhile, TOC loss was highest in light exposed HF treatments (i.e., HF and HF+RO) and the HF samples had an upward trend in BP throughout the experiment with BP of both, HF and HF+RO samples, being significantly higher at the end of the experiment compared to the other treatments. This suggests that exposure to light reduced the inhibitory effect of HF, and made it prone to mineralization, likely via bacterial degradation and photomineralization. Although not specifically tested in this experiment, such process could occur in treatments with other leonardite products, e.g. SH in a similar way.

The importance of light in determining the fate of the different agents utilized in our browning experiments was corroborated by the results of both the DOM quality assessed by EEMs and the BCC. The EEM comparison of the dark and light incubations of the microbial and alkalinity tests clearly demonstrated that the major qualitative changes in DOM depended on the exposure to light, and these changes were especially related to high molecular weight substances and substances related to biological activity. Furthermore, although all browning agents were impacted by light, it affected HF and SH differently from RO. As changes in fluorescence peaks in light were the smallest for the soft water samples, a possible explanation of the impact of light could be the fact that EEM spectroscopy probes only the quality of DOC, and not POC. Hence, EEM changes may at least in part reflect the transformation of DOC to POC due to flocculation. However, one of the most substantial differences in EEM profiles between light and dark incubations were detected in the HF samples of the microbial test where POC formation was unaffected by light. This further supports the idea that light exposure may have initiated substantial photochemical changes in the leonardite agents other than flocculation. In support of this hypothesis, we observed significant loss in color in the HF treatments but not in the RO treatments (Figure 1a). These results agree with Lennon, Hamilton, Muscarella and others (2013), who proposed that the leonardite product SH may be more prone to the loss of color than DOC found in natural lakes.

### Biotic effects: bacterial responses

The light-induced change in DOM also affected the BCC. The structure of the bacterial communities that developed in the HF treatments (i.e., HF and HF+RO treatments of the mesocosms and light test) clearly showed that this agent modified the bacterial communities when exposed to light. The lack of these specific differences in dark incubations suggest that the exposure to light is necessary for HF to become available for utilization by bacterial communities. This idea is supported by the results of the BP measurements as well as the results of the TOC changes during the same experiment. Combined these results suggest that light exposure increased the bioavailability of HF for heterotrophic bacteria and enhanced the mineralization of this organic carbon pool. Such potential mineralization of HF upon exposure to light substantially compromises the concept of its use as an inert “sunscreen” in browning experiments.

### Biotic effects: zooplankton life history responses

In the reproduction experiment, we could not detect any stimulation of *Daphnia* growth by any of the tested browning agents. We therefore conclude that neither HF, SH nor RO would act as a subsidy for zooplankton. On the contrary, all three browning agents had negative impacts on the zooplankton. The strongest impact was seen for HF, which in the mesocosms, reduced the abundance of Cladocera and Copepoda on average by the factor of five and nine, respectively, and had negative impacts on the reproduction of daphnids. One potential reason for the negative effects on Cladocera in the mesocosms could be the higher rates of particle formation observed in treatments with HF, which could interfere with the filter-feeding apparatus of the Cladocera, potentially compromising their feeding and digestion. However, this would not explain the detrimental effects on Copepoda observed in the mesocosms, as these are raptorial feeders (Brandl 2005). Another potential reason for the fitness impairments of HF on zooplankton could come from stress responses. In a previous study, *D. magna* responded to HF treatments with an increase in antioxidant capacity and oxidative damage, combined with a reduced amount of energy available (Saebelfeld, Minguez, Griebel and others 2017).

Despite the strong impact of HF on the zooplankton in the mesocosms, it did not have a significant impact in the acute immobilization test of cultured *D. magna*, although similar concentrations were used. Possibly, *D. magna*, as one of the biggest freshwater Cladocera species, is more tolerant against harmful substances compared to the smaller species present in a natural cladoceran community (Koivisto 1995; Saebelfeld, Minguez, Griebel and others 2017). Still, all browning agents had a negative impact on the average number of offspring per clutch in the reproduction experiment. The total number of offspring was significantly lower compared to the control only in HF, but there was a decreasing trend also for SH and RO, suggesting that those may also affect *Daphnia* reproduction. These results are in line with a previous study that reported delayed maturity and reduced number of offspring combined with stress induction in *D. magna* in experiments using HF at a slightly higher concentration (Saebelfeld, Minguez, Griebel and others 2017). This study further reported no offspring production in *D. longispina* at high concentrations of HF (30 mg DOC L ^-1^). Furthermore, Bouchnak and Steinberg (2013) saw a similar trend of decreased egg production in *D. magna* in the presence of HF, even at a concentration of 5 mg C L ^-1^, but at the same time they also found that the lifespan of the Cladocera increased. Nevertheless, there are also some earlier findings that contradict the negative effects of browning agents on zooplankton observed in our experiments. In a study testing SH, Lennon, Hamilton, Muscarella and others (2013) did not find any evidence of negative effects of SH on fitness of *D. pulex x pulicaria* clones, but in fact, reported a 10 % increase in the intrinsic rate of increase in a life table experiment conducted with *Daphnia*. However, they note that the positive impact was marginal and that additional experiments are needed. Our combined results of the zooplankton testing suggest dramatic and negative effects of HF on the zooplankton community. This could result in strong direct and indirect effects on overall ecosystem processes with implications also for the fish and phytoplankton. Nonetheless, the underlying causes for the adverse effects of HF on zooplankton still remain elusive.

### Biotic effects: G. semen growth dynamics responses

The effect of all three browning agents on the growth rate of *G. semen* were tested, but none of them had, not even at high concentrations, any negative effects. Overall, these results indicate high tolerance of *G. semen* to all three substances. Not only does this demonstrate a lack of toxic effects, but also that no light limitation was generated in these experiments. Sassenhagen, Wilken, Godhe and others (2015) showed that *G. semen* growth rates were only slightly reduced when light intensity was dropped from 150 to 25 µmol photons m^-2^ s^-1^. In a mesocosm or alike, it is possible that the different browning agents could have a larger effect. Growth rates were higher in treatments with RO compared to the control. Unfortunately, the RO agents were contaminated with a green algae, demonstrating that the risk of contamination using RO can be considered high, and can be explained by its origin from an active aquatic environment. RO at intermediate concentrations furthermore showed the highest variation among replicates. Therefore, extra precautions must be taken when utilizing this agent.

## Conclusion

The purpose of this study was to evaluate the use of commercially available leonardite browning agents as an experimental analogue for indigenous DOC of terrestrial origin. Compared to RO, HF and SH are biogeochemically highly distinct. These compounds are primarily used in experiments to mimic browning of natural waters, but our results showed that the water color-modifying effect of the leonardite products decreases gradually upon exposure to light. Our results further suggest that besides the loss of color, light exposure also prompts other changes in DOM quality that lead to enhanced mineralization of organic carbon, and alterations in the composition of bacterial communities. All light-induced changes substantially compromise the concept of using leonardite in browning experiments where it is expected to act as a practically inert, recalcitrant chromophore. Moreover, having chemical properties substantially different from the DOC associated with natural browning, it is also a questionable analogue of terrestrial DOC as a subsidy to aquatic ecosystems. However, as the severity of changes was related to water chemistry (e.g., alkalinity), even leonardite may be an acceptable alternative in some cases. For example, the extent of particle formation and the quality changes of DOM depended on the alkalinity of water with substantially fewer negative effects in soft than in hard water. Further, all tested agents had negative impacts on zooplankton, with the most severe seen for HF. The impairments may be due to stress induction, but the exact mechanisms should be further investigated. Another specific problem that could arise from the use of DOM concentrates extracted from active ecosystems is the unintentional addition of local biota that as a biological contamination might bias the results of manipulation experiments, as seen from the green algae contamination in RO treatment of *G. semen* cultures.

In conclusion, our extensive tests of leonardite products raise multiple concerns on their suitability as proxies for natural browning of freshwater ecosystems. Our experiments (this study, Attermeyer, Andersson, Catalán and others 2019; Nydahl, Wallin, Tranvik and others 2019, Chaguaceda and others in preparation) show that it is feasible to prepare reverse osmosis concentrates for browning of several thousand litres. Hence, we recommend, 1) that browning agents derived from humic aquatic environments or soils, such as RO concentrates, should be prioritized at the laboratory scale and for mesocosms with careful consideration of potential biotic contaminations, and 2) where leonardite extracts are used, great attention should be paid to effects that may be atypical for indigenous browning agents.

## Acknowledgements

Financial support was received from the Knut and Alice Wallenberg Foundation for funding to LJT (Grant KAW 2013.0091). We are extremely grateful to all those people involved in maintaining and sampling the mesocosms, especially William Colom Montero, Fernando Chaguaceda, Sara Andersson, Carolin Hiller, and Holger Villwock for all the help in the field and the laboratory. Thanks also to Don Pierson, Björn Mattson, and Christer Strandberg for technical assistance and to Jean Pettersson, Helena Enderskog, Kristiina Mustonen and Erika Bridell for help with laboratory analyses. Mafalda Castro gave valuable insight and guidance in *Daphnia* culturing. Judita Koreivienė provided the *G. semen* monoculture.

