## Supplementary Material for "Comprehensive analysis of chemical and biological problems associated with browning agents used in aquatic studies"

### Appendix 1:

*Technical parameters of the chemical characterization of browning agents*

*(a) Chemical characterization*

Samples were analyzed after digestion of the solid (HuminFeed (HF) and SuperHume (SH)) or liquid (Reverse Osmosis (RO)) sample (0.2 g sample in 50 ml micro Kjedahl flask). To each sample, 3 ml nitric acid and 1.5 ml hydrogen peroxide were added and heated to 80°C overnight and thereafter at 145°C for 6-8 hour until an almost clear (slightly yellow) solution appeared. The samples were diluted to 40.0 mL in the Kjeldahl flasks. On the Spectro Ciros CCD ICP-AES (Spectro, Kleve, Germany) used for the determination of trace elements in the digested samples for all elemental wavelengths, proper background subtractions were performed and the net-signals were used for further calculations. The samples were aspirated at 2 ml min $^{-1}$ to a modified Lichte nebulizer with a cyclonic spray-chamber. The argon flow rates were 0.9 L min $^{-1}$ for nebulization, 0.9 L min $^{-1}$ for the auxiliary gas supply and 14.0 L min $^{-1}$ as coolant gas. The plasma power was set to 1400 W at 27 MHz. The samples were aspirated for 45 s before three 24 s integrations of the chosen lines were performed. The average signal was used for further calculations, and calibration was performed against matrix matched certified reference solutions (Spectrascan, Teknolab AS, Norway).

*(b) Nuclear Magnetic Resonance (NMR)*

$^{1}$H NMR was conducted on a Bruker advanced Neo 600 MHz spectrometer (1H NMR: 600.18 MHz), equipped with a cryogenic tippled resonance probe TCI (CRPHe TR-1H &19F/13C/15N 5mm-EZ). A pulse program (from the manufacturer) designed to suppress solvent signals by excitation sculpting was used. The middle of the spectrum was set on-resonance to the solvent signals. A total of 1024 scans with 32K points and a relaxation delay of 2 seconds were acquired for each experimental sample. All NMR spectra were processed and analyzed with the Bruker Topspin and MNova software programs. Chemical shifts are reported in parts per million (ppm) on the δ scale from an internal standard.

*(c) High Pressure Size Exclusion Chromatography – High Resolution Mass Spectrometry (HPSEC-HRMS)*

HPSEC- HRMS was conducted with an Agilent 1100 HPLC (Agilent, Santa Clara, CA, USA) equipped with a UV-Vis Diode Array Detector for sample light attenuation (Agilent 1100, Santa Clara, CA, USA) and an Orbitrap mass spectrometer (LTQ-Velos Pro, Thermo Fisher Scientific, Waltham, MA, USA) in series that detected negatively ioniziable molecules by electrospray ionization mass spectrometry, as described in Hawkes, Sjöberg, Bergquist and others (2019). 20 μL of sample was injected onto a size exclusion column (Tosoh TSK Gel G3000SW, 300 x 7.5 mm, 10 μm), and the sample was eluted with 25 mM ammonium acetate dissolved in 20:80 methanol:water at 1 ml min $^{-1}$. The Orbitrap was tuned for maximum signal at 369.1 Da with SRFA, and data were collected at a spray voltage of 2.5 kV in negative mode, measuring between 200-2000 Da.

**Appendix 2**: $^{1}$H NMR chemical shifts. Blue = SRFA, Green = HF, Red = SH. 0-1.6 ppm = aliphatic terpenoids, 1.6-3.2 ppm = carboxylic rich alicyclic material, 3.2-4.3 ppm= carbohydrates, 6-8.5= aromatic compounds. The peaks at 1.8, 2.0, 3.2, and 8.3 ppm are solvent peaks.

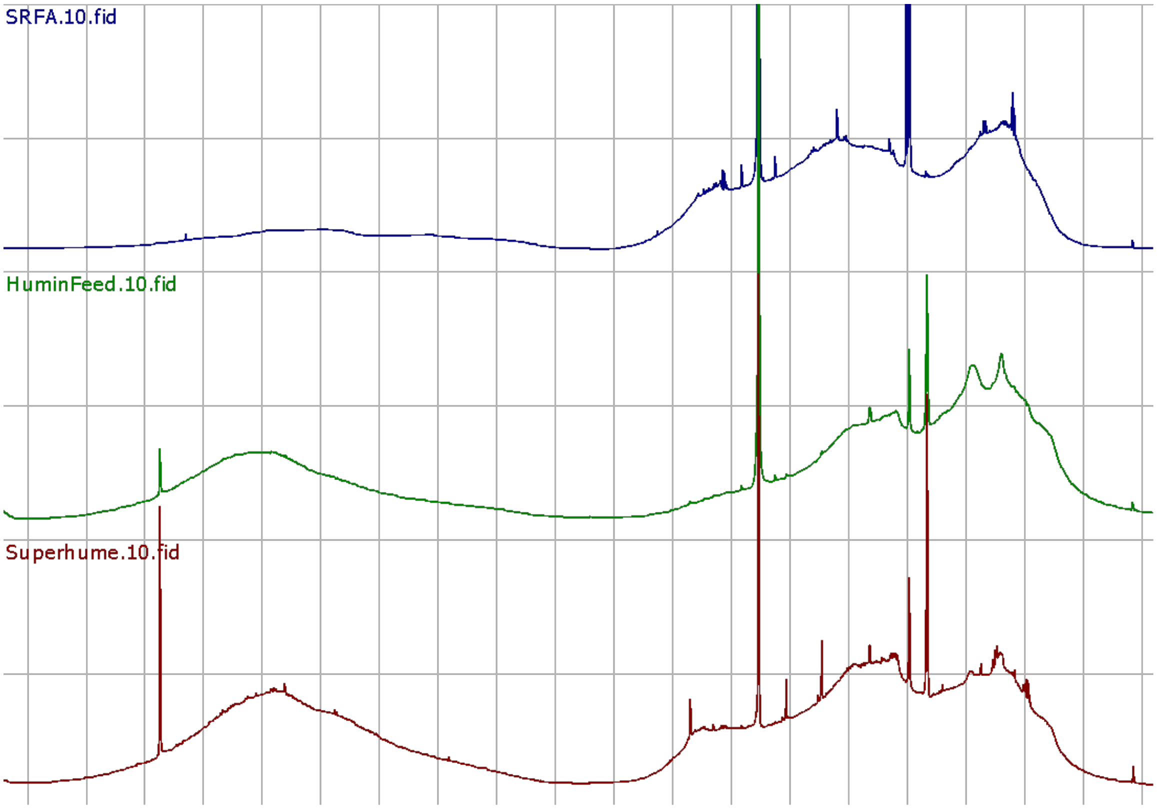

**Appendix 3**: High Pressure Size Exclusion Chromatography - High Resolution Mass Spectrometry (HPSEC-HRMS) of HF, SH and RO (here SRFA). Blue lines depict absorbance at 420 nm, whereas orange lines depict ionized material with *m/z* 350-450 measured by electrospray ionization mass spectrometry.

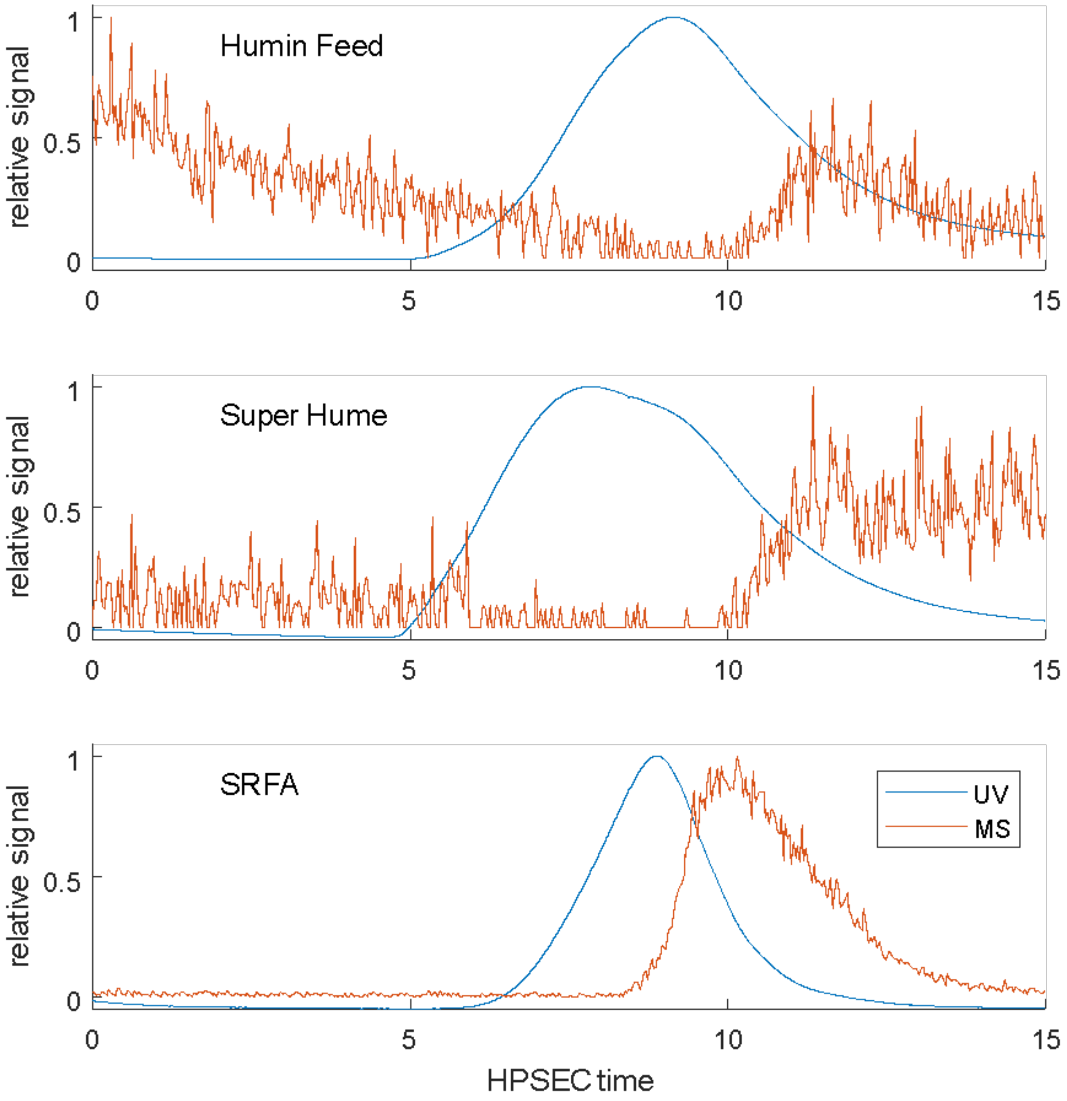

**Appendix 4:** *P*-values of pairwise comparisons of browning agents from a *post-hoc* test on flocculation occurring in the mesocosms. Bold fonts depict significant differences.

| | **Control** | **HF** | **HF+RO** | **RO** |
|---|---|---|---|---|
| **Control** | 1.000 | **<0.001** | **<0.001** | 0.222 |
| **HF** | | 1.000 | 0.999 | **<0.001** |
| **HF+RO** | | | 1.000 | **<0.001** |
| **RO** | | | | 1.000 |

**Appendix 5:** Changes in water color measured as $Abs_{420}$ during the microbial test for samples incubated in dark and light with different browning agents (Control, HF and HF+RO and RO).

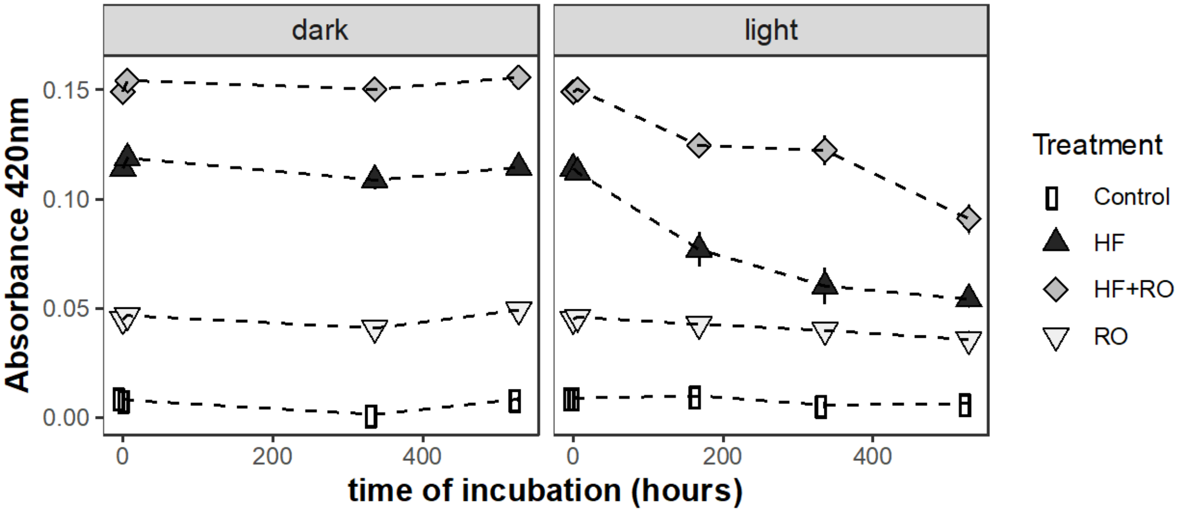

**Appendix 6**: P-values of pairwise comparisons of browning agents from the *post-hoc* tests on flocculation (measured as POC, TOC, or particle mass (PM)) under dark or light conditions occurring in the a) microbial and b) alkalinity test. Bold fonts depict significant differences.

***a) Microbial test***

| **Parameter** | | Control – dark | HF – dark | HF+RO – dark | RO – dark | Control – light | HF – light | HF+RO – light | RO – light |
|---|---|---|---|---|---|---|---|---|---|
| **I) $Abs_{420}$** | Control – dark | 1.000 | 0.999 | **0.048** | 0.397 | 0.981 | **<0.001** | **<0.001** | **0.010** |
| | HF – dark | | 1.000 | 0.124 | 0.675 | 0.859 | **<0.001** | **<0.001** | **0.003** |
| | HF+RO – dark | | | 1.000 | 0.939 | **0.006** | **<0.001** | **<0.001** | **<0.001** |
| | RO – dark | | | | 1.000 | 0.079 | **<0.001** | **<0.001** | **<0.001** |
| | Control – light | | | | | 1.000 | **<0.001** | **<0.001** | 0.079 |
| | HF – light | | | | | | 1.000 | 0.991 | **<0.001** |

| | | | | | | | | | |
|---|---|---|---|---|---|---|---|---|---|
| | HF+RO – light | | | | | | | 1.000 | **<0.001** |
| | RO – light | | | | | | | | 1.000 |
| **II) log POC** | Control – dark | 1.000 | **<0.001** | **<0.001** | 0.103 | **<0.001** | **<0.001** | **<0.001** | **<0.001** |
| | HF – dark | | 1.000 | 0.999 | **0.010** | 0.996 | 0.940 | 0.075 | 0.296 |
| | HF+RO – dark | | | 1.000 | **0.003** | 0.902 | 0.999 | 0.224 | 0.625 |
| | RO – dark | | | | 1.000 | **0.049** | **0.001** | **<0.001** | **<0.001** |
| | Control – light | | | | | 1.000 | 0.605 | **0.016** | 0.083 |
| | HF – light | | | | | | 1.000 | 0.514 | 0.913 |
| | HF+RO – light | | | | | | | 1.000 | 0.994 |
| | RO – light | | | | | | | | 1.000 |

| | | | | | | | | | |
|---|---|---|---|---|---|---|---|---|---|
| **III) log DOC** | Control – dark | 1.000 | 1.000 | **<0.001** | **<0.001** | **0.001** | **<0.001** | **<0.001** | **<0.001** |
| | HF – dark | | 1.000 | **<0.001** | **0.001** | **<0.001** | **<0.001** | **<0.001** | **<0.001** |
| | HF+RO – dark | | | 1.000 | 0.645 | **<0.001** | **<0.001** | **<0.001** | **0.001** |
| | RO – dark | | | | 1.000 | **<0.001** | **0.009** | **<0.001** | 0.091 |
| | Control – light | | | | | 1.000 | **<0.001** | **<0.001** | **<0.001** |
| | HF – light | | | | | | 1.000 | 0.096 | 0.960 |
| | HF+RO – light | | | | | | | 1.000 | **0.009** |
| | RO – light | | | | | | | | 1.000 |
| **IV) TOC** | Control – dark | 1.000 | 0.745 | 0.990 | **0.028** | 0.056 | **<0.001** | **<0.001** | **0.016** |
| | HF – dark | | 1.000 | 0.274 | **0.001** | 0.714 | **<0.001** | **<0.001** | **<0.001** |

| | | | | | | | | | |
|---|---|---|---|---|---|---|---|---|---|
| | HF+RO – dark | | | 1.000 | 0.154 | **0.009** | **<0.001** | **<0.001** | 0.097 |
| | RO – dark | | | | 1.000 | **<0.001** | **<0.001** | **<0.001** | 1.000 |
| | Control – light | | | | | 1.000 | **<0.001** | **<0.001** | **<0.001** |
| | HF – light | | | | | | 1.000 | **<0.001** | **<0.001** |
| | HF+RO – light | | | | | | | 1.000 | **<0.001** |
| | RO – light | | | | | | | | 1.000 |

***b) Alkalinity test***

| **Parameter** | | Control – dark | HF – dark | SH – dark | RO – dark | Control – light | HF – light | SH – light | RO – light |
|---|---|---|---|---|---|---|---|---|---|
| **I) PM hard water** | Control – dark | 1.000 | **<0.001** | **<0.001** | 0.912 | 1.000 | **<0.001** | **<0.001** | 0.252 |

| | | | | | | | | | |
|---|---|---|---|---|---|---|---|---|---|
| | HF – dark | | 1.000 | 0.995 | **<0.001** | **<0.001** | 0.088 | 0.414 | **0.010** |
| | SH – dark | | | 1.000 | **<0.001** | **<0.001** | 0.227 | 0.756 | **0.004** |
| | RO – dark | | | | 1.000 | 0.820 | **<0.001** | **<0.001** | 0.799 |
| | Control – light | | | | | 1.000 | **<0.001** | **<0.001** | 0.217 |
| | HF – light | | | | | | 1.000 | 0.969 | **<0.001** |
| | SH – light | | | | | | | 1.000 | **0.001** |
| | RO – light | | | | | | | | 1.000 |
| **II) PM soft water** | Control – dark | 1.000 | 0.799 | 0.999 | 1.000 | 1.000 | 0.067 | 0.999 | 1.000 |

| | | | | | | | | | |
|---|---|---|---|---|---|---|---|---|---|
| | HF – dark | | 1.000 | 0.972 | 0.895 | 0.715 | 0.484 | 0.968 | 0.970 |
| | SH – dark | | | 1.000 | 1.000 | 0.991 | 0.147 | 1.000 | 1.000 |
| | RO – dark | | | | 1.000 | 0.999 | 0.094 | 1.000 | 1.000 |
| | Control – light | | | | | 1.000 | 0.068 | 0.992 | 0.998 |
| | HF – light | | | | | | 1.000 | 0.141 | 0.184 |
| | SH – light | | | | | | | 1.000 | 1.000 |
| | RO – light | | | | | | | | 1.000 |

**Appendix 7:** Comparisons of initial and final TOC concentrations in the different treatments (i.e Control, HF, HF+RO) of the microbial test.

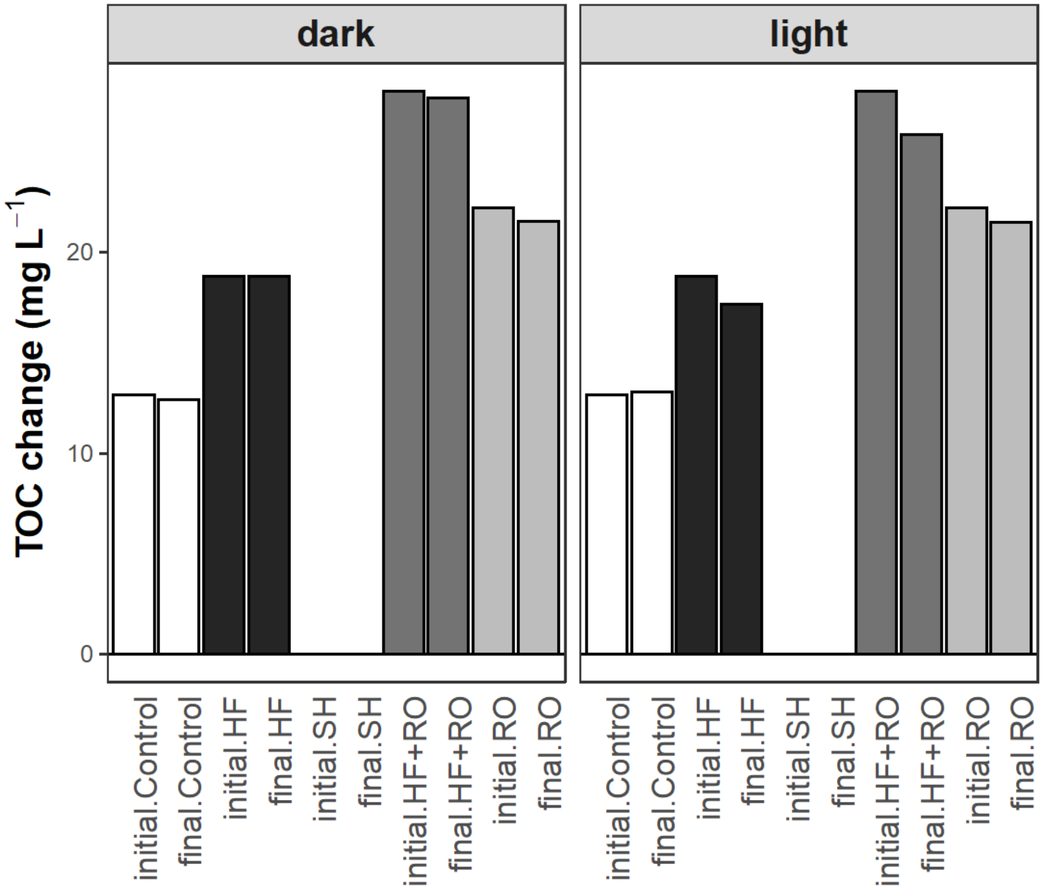

**Appendix 8**: Change of DOC concentrations in the different treatments (i.e. Control, HF, RO, HF+RO) over the course of the microbial experiment.

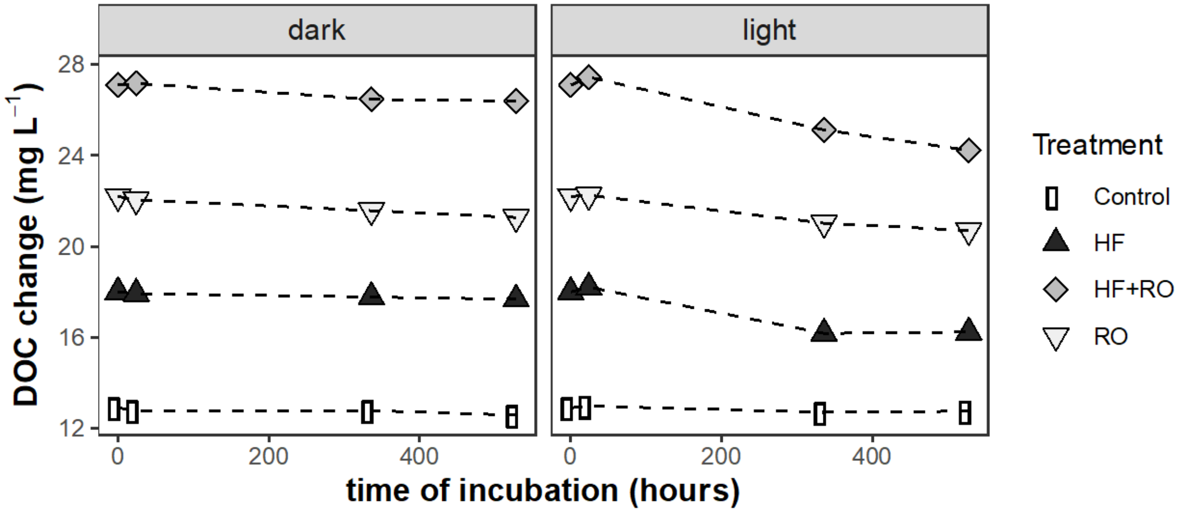

**Appendix 9**: $R^2$ and *p*-values of PERMANOVA analyses (*adonis* function) of multivariate data of EEM spectroscopy changes, and the bacterial community composition (BCC) of the mesocosms and the microbial test using the factors light (levels: dark or light), hardness (levels: soft or hard) and agent (levels: control, RO and leonardite containing agent (i.e., HF, HF+RO, SH). Bold fonts depict significant differences. df: degrees of freedom.

| **Factor (df)** | **EEMS** | | **BCC mesocosms** | | **BCC microbial test** | |
|---|---|---|---|---|---|---|
| | $R^2$ | *p* | $R^2$ | *p* | $R^2$ | *p* |
| Light (1) | -0.82147 | 0.992 | | | 0.31262 | **0.001** |
| Hardness (1) | 1.01993 | **0.002** | | | | |
| Agent (2) | 0.89624 | **0.007** | 0.60528 | **0.001** | 0.21046 | **0.001** |
| Light * hardness (1) | 0.79635 | **0.006** | | | | |
| Light * agent (2) | 0.7024 | **0.027** | | | 0.18942 | **0.001** |
| Hardness * agent (2) | -0.91478 | 0.996 | | | | |
| Light * hardness * agent (2) | -0.67887 | 0.991 | | | | |

**Appendix 10**: Changes in extracted EEM spectroscopy peaks.

**(a) Changes in peak A related to high molecular weight and aromatic humic substances**

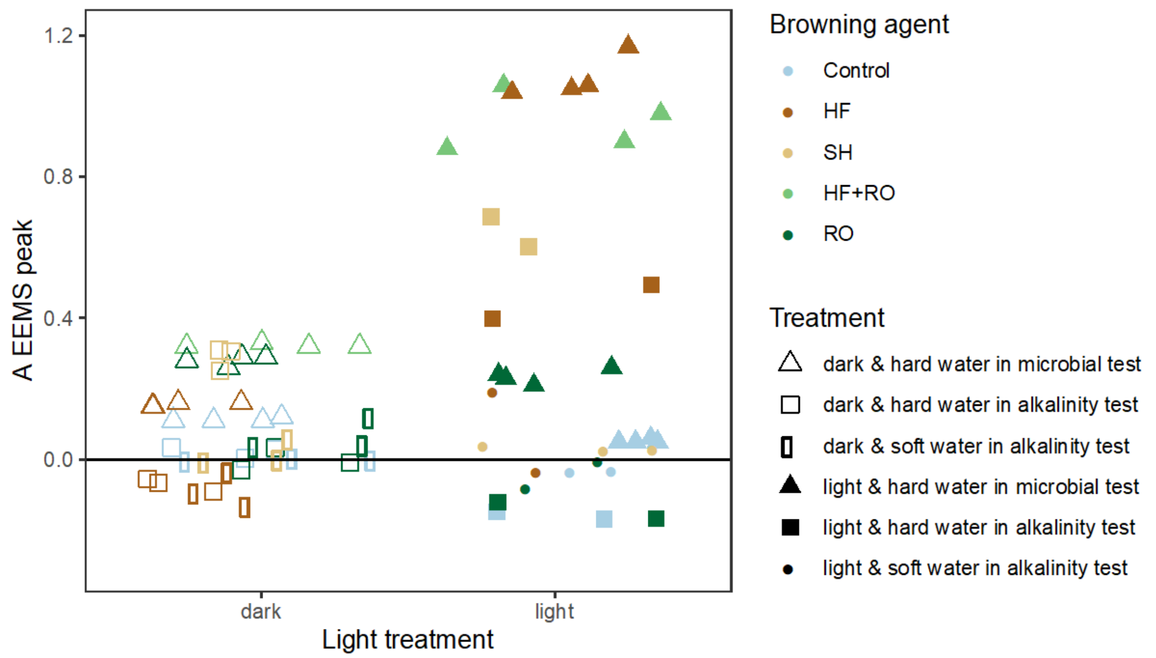


**(b) Changes in peak M related to low molecular weight substances with biological activity**

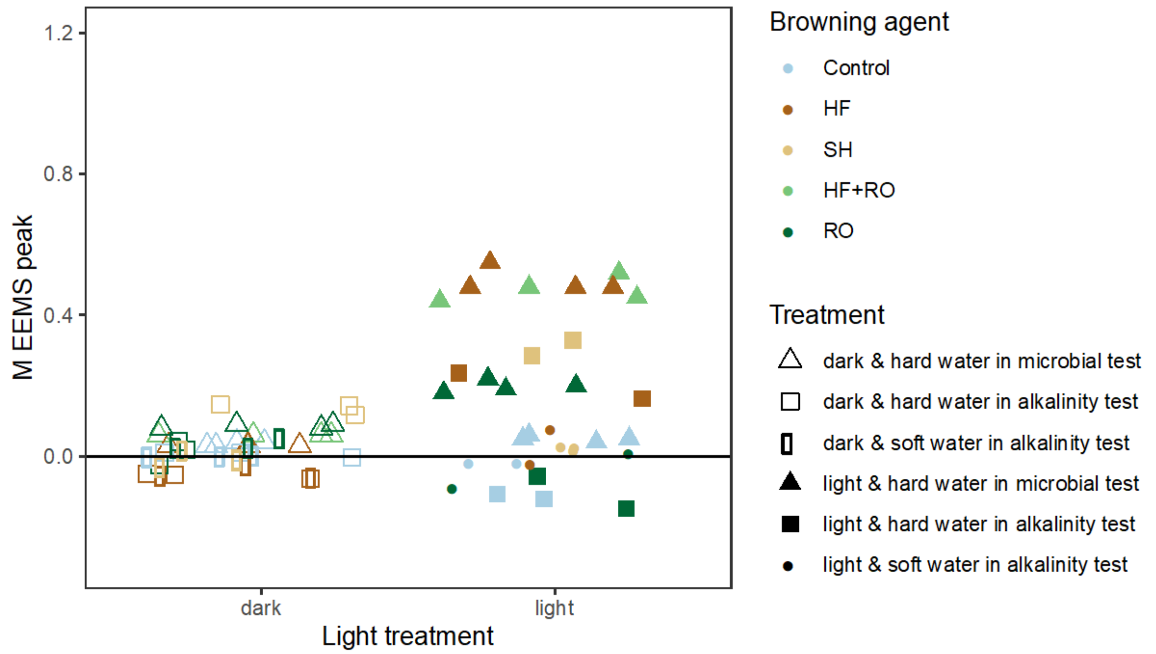

**(c) Changes in peak T related tryptophan, indicating intact or less degraded proteins**

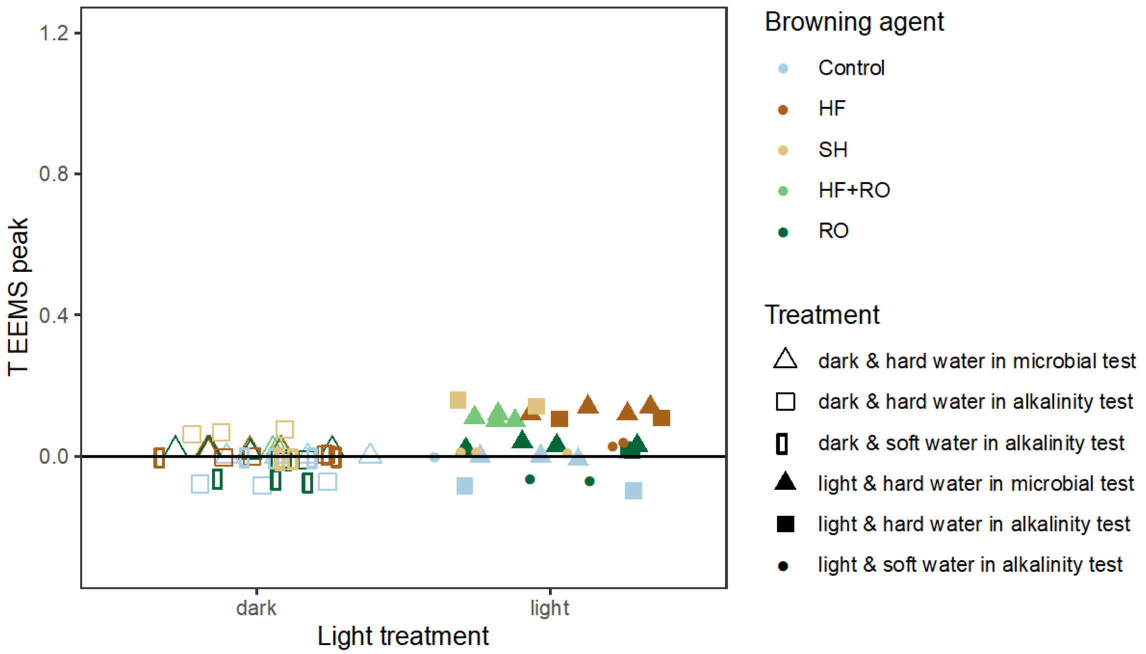


**(d) Changes in peak C related to high molecular weight humic substances**

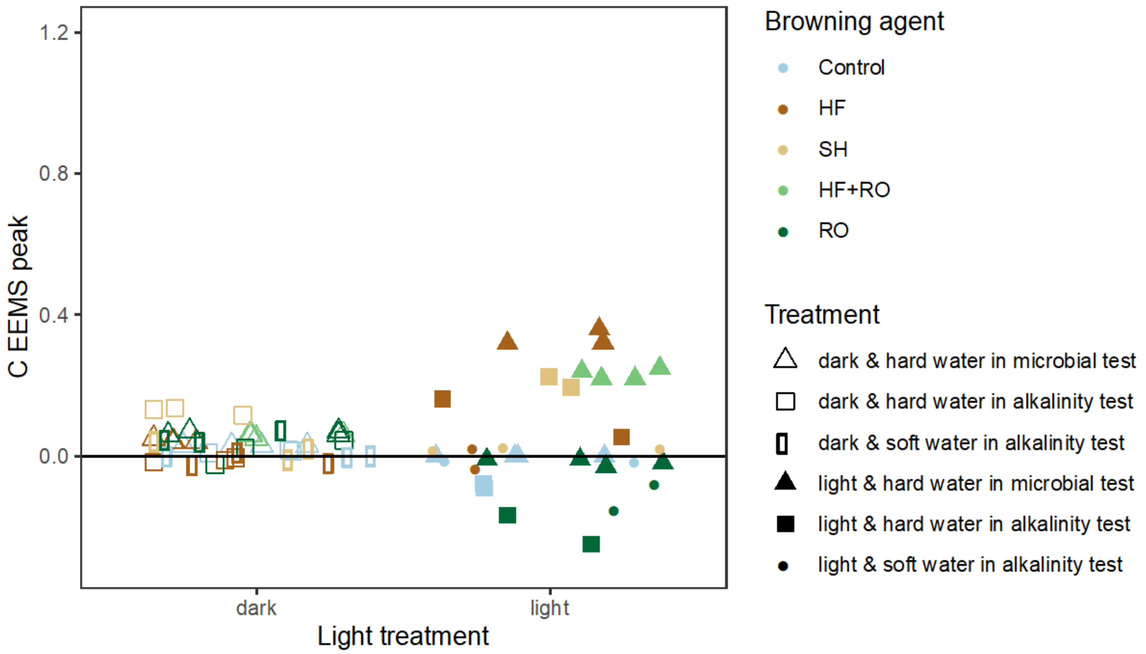

**(e) Changes in peak B related to tyrosine, indicating degraded proteins**

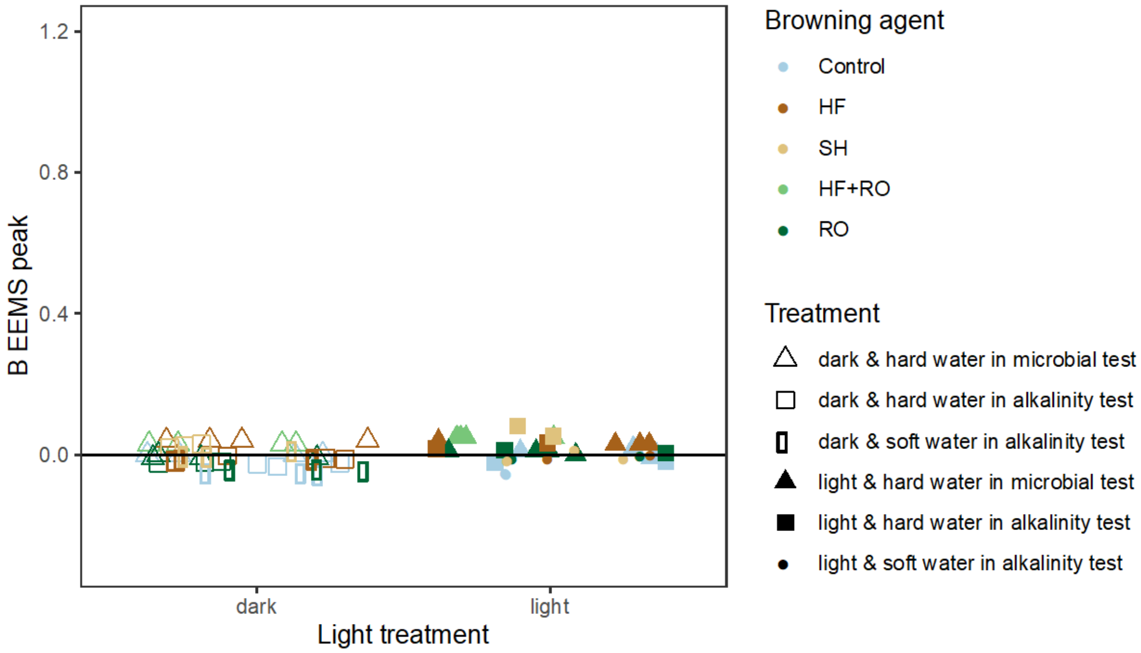

**Appendix 11:** Relative abundance of major bacterial families at the end of the microbial test

**(f) Dark incubation** **(b) Light incubation**

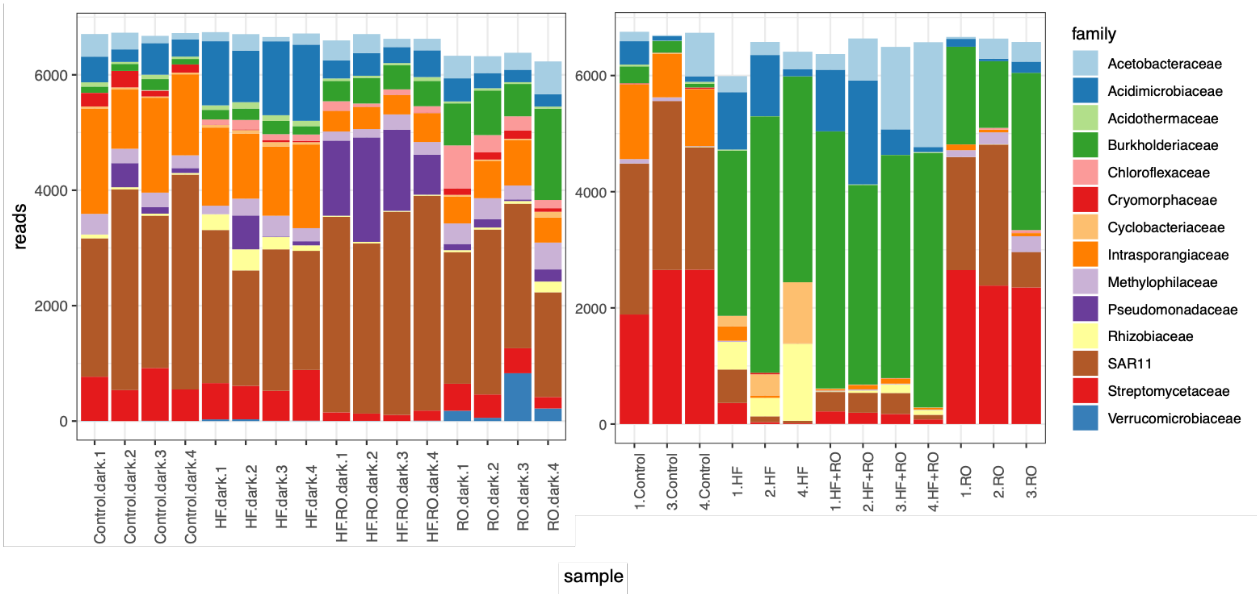

**Appendix 12**: *P*-values of pairwise *post-hoc* tests of the effect of browning agents on bacterial production at the end of the microbial test under dark (a) and light (b) incubation. Bold fonts depict significant differences.

**a) Dark incubation**

| | **Control** | **HF** | **HF+RO** | **RO** |
|---|---|---|---|---|
| **Control** | 1.000 | 0.838 | **<0.001** | **<0.001** |
| **HF** | | 1.000 | **0.002** | **<0.001** |
| **HF+RO** | | | 1.000 | 0.894 |
| **RO** | | | | 1.000 |

**b) Light incubation**

| | **Control** | **HF** | **HF+RO** | **RO** |
|---|---|---|---|---|
| **Control** | 1.000 | **0.002** | **<0.001** | 0.274 |
| **HF** | | 1.000 | 0.852 | **0.046** |
| **HF+RO** | | | 1.000 | **0.011** |
| **RO** | | | | 1.000 |

**Appendix 13**: Coefficients of linear model depicting pairwise comparison of a) zooplankton abundance between mesocosms treated with different browning agents, b) number of offspring per clutch, and c) total number of *D. magna* offspring during a 21-day reproduction test with different browning agents. Bold fonts depict significant results. df: degrees of freedom.

| Measure | Group | Comparison | df | *t*-value | *P*-value |
|---|---|---|---|---|---|
| **a) Zooplankton abundance** | I) Cladocera | **Control -HF** | **3,16** | **-2.485** | **0.0244** |
| | | **Control – HF+RO** | **3,16** | **-3.170** | **0.0059** |
| | | Control – RO | 3,16 | -1.069 | 0.3010 |
| | II) Copepoda | **Control -HF** | **3,16** | **-2.775** | **0.0135** |
| | | **Control - HF+RO** | **3,16** | **-4.675** | **<0.001** |
| | | Control – RO | 3,16 | 0.513 | 0.6152 |
| **c) Number of offspring per clutch** | *D. magna* | **Control -HF** | **3, 25** | **-5.108** | **<0.001** |
| | | **Control – SH** | **3, 25** | **-2.768** | **0.010** |
| | | **Control – RO** | **3, 25** | **-2.772** | **0.010** |
| **b) Total number of offspring** | *D. magna* | **Control -HF** | **3, 25** | **-3.499** | **0.0018** |
| | | Control – SH | 3, 25 | -1.439 | 0.0549 |
| | | Control – RO | 3, 25 | -2.014 | 0.1479 |

**Appendix 14**: *P*-values of pairwise comparisons of browning agents of different concentrations from a *post-hoc* test on growth rates of *Gonyostomum semen*. Bold fonts depict significant differences.

| | | **Control** | **HF** | | | **SH** | | | **RO** | |
|---|---|---|---|---|---|---|---|---|---|---|
| | | | **low** | **medium** | **high** | **low** | **medium** | **high** | **low** | **medium** |
| **Control** | | 1.000 | 0.986 | 1.000 | 0.256 | 0.942 | 0.996 | 1.000 | 0.454 | **0.003** |
| **HF** | **low** | | 1.000 | 0.990 | **0.031** | 0.422 | 0.698 | 0.995 | 0.954 | **0.036** |
| | **medium** | | | 1.000 | 0.236 | 0.929 | 0.994 | 1.000 | 0.483 | **0.003** |
| | **high** | | | | 1.000 | 0.930 | 0.733 | 0.198 | **0.001** | **<0.001** |
| **SH** | **low** | | | | | 1.000 | 1.000 | 0.898 | **0.039** | **<0.001** |
| | **medium** | | | | | | 1.000 | 0.988 | 0.108 | **<0.001** |
| | **high** | | | | | | | 1.000 | 0.544 | **0.004** |
| **RO** | **low** | | | | | | | | 1.000 | 0.399 |
| | **medium** | | | | | | | | | 1.000 |
